# A chromosome region linked to neurodevelopmental disorders acts in distinct neuronal circuits in males and females to control locomotor behavior

**DOI:** 10.1101/2024.05.17.594746

**Authors:** Jaekyoon Kim, Yann Vanrobaeys, M. Felicia Davatolhagh, Benjamin Kelvington, Snehajyoti Chatterjee, Sarah L. Ferri, Christopher Angelakos, Alea A. Mills, Marc V. Fuccillo, Thomas Nickl-Jockschat, Ted Abel

**Affiliations:** Department of Neuroscience and Pharmacology, Carver College of Medicine, University of Iowa, IA; Iowa Neuroscience Institute, University of Iowa, IA; Interdisciplinary Graduate Program in Genetics, University of Iowa, IA; Department of Neurobiology, David Geffen School of Medicine, University of California Los Angeles, CA; Stead Family Department of Pediatrics, Carver College of Medicine, University of Iowa, IA; Department of Psychiatry and Behavioral Sciences, Stanford University School of Medicine, Stanford, CA; Cold Spring Harbor Laboratory, Cold Spring Harbor, NY; Department of Neuroscience, University of Pennsylvania, PA; Department of Psychiatry and Psychotherapy, Otto von Guericke University Magdeburg, Germany; German Center for Mental Health (DZPG), partner site Halle-Jena-Magdeburg, Germany; Center for Intervention and Research on adaptive and maladaptive brain Circuits underlying mental health (C-I-R-C), Halle-Jena-Magdeburg, Germany; Department of Psychiatry, Carver College of Medicine, University of Iowa, IA

## Abstract

Biological sex shapes the manifestation and progression of neurodevelopmental disorders (NDDs). These disorders often demonstrate male-specific vulnerabilities; however, the identification of underlying mechanisms remains a significant challenge in the field. Hemideletion of the 16p11.2 region (16p11.2 del/+) is associated with NDDs, and mice modeling 16p11.2 del/+ exhibit sex-specific striatum-related phenotypes relevant to NDDs. Striatal circuits, crucial for locomotor control, consist of two distinct pathways: the direct and indirect pathways originating from D1 dopamine receptor (D1R) and D2 dopamine receptor (D2R) expressing spiny projection neurons (SPNs), respectively. In this study, we define the impact of 16p11.2 del/+ on striatal circuits in male and female mice. Using snRNA-seq, we identify sex- and cell type-specific transcriptomic changes in the D1- and D2-SPNs of 16p11.2 del/+ mice, indicating distinct transcriptomic signatures in D1-SPNs and D2-SPNs in males and females, with a ∼5-fold greater impact in males. Further pathway analysis reveals differential gene expression changes in 16p11.2 del/+ male mice linked to synaptic plasticity in D1- and D2-SPNs and GABA signaling pathway changes in D1-SPNs. Consistent with our snRNA-seq study revealing changes in GABA signaling pathways, we observe distinct changes in miniature inhibitory postsynaptic currents (mIPSCs) in D1- and D2-SPNs from 16p11.2 del/+ male mice. Behaviorally, we utilize conditional genetic approaches to introduce the hemideletion selectively in either D1- or D2-SPNs and find that conditional hemideletion of genes in the 16p11.2 region in D2-SPNs causes hyperactivity in male mice, but hemideletion in D1-SPNs does not. Within the striatum, hemideletion of genes in D2-SPNs in the dorsal lateral striatum leads to hyperactivity in males, demonstrating the importance of this striatal region. Interestingly, conditional 16p11.2 del/+ within the cortex drives hyperactivity in both sexes. Our work reveals that a locus linked to NDDs acts in different striatal circuits, selectively impacting behavior in a sex- and cell type-specific manner, providing new insight into male vulnerability for NDDs.

**Highlights:**

- 16p11.2 hemideletion (16p11.2 del/+) induces sex- and cell type-specific transcriptomic signatures in spiny projection neurons (SPNs).
- Transcriptomic changes in GABA signaling in D1-SPNs align with changes in inhibitory synapse function.
- 16p11.2 del/+ in D2-SPNs causes hyperactivity in males but not females.
- 16p11.2 del/+ in D2-SPNs in the dorsal lateral striatum drives hyperactivity in males.
- 16p11.2 del/+ in cortex drives hyperactivity in both sexes.

**Graphic abstract:** 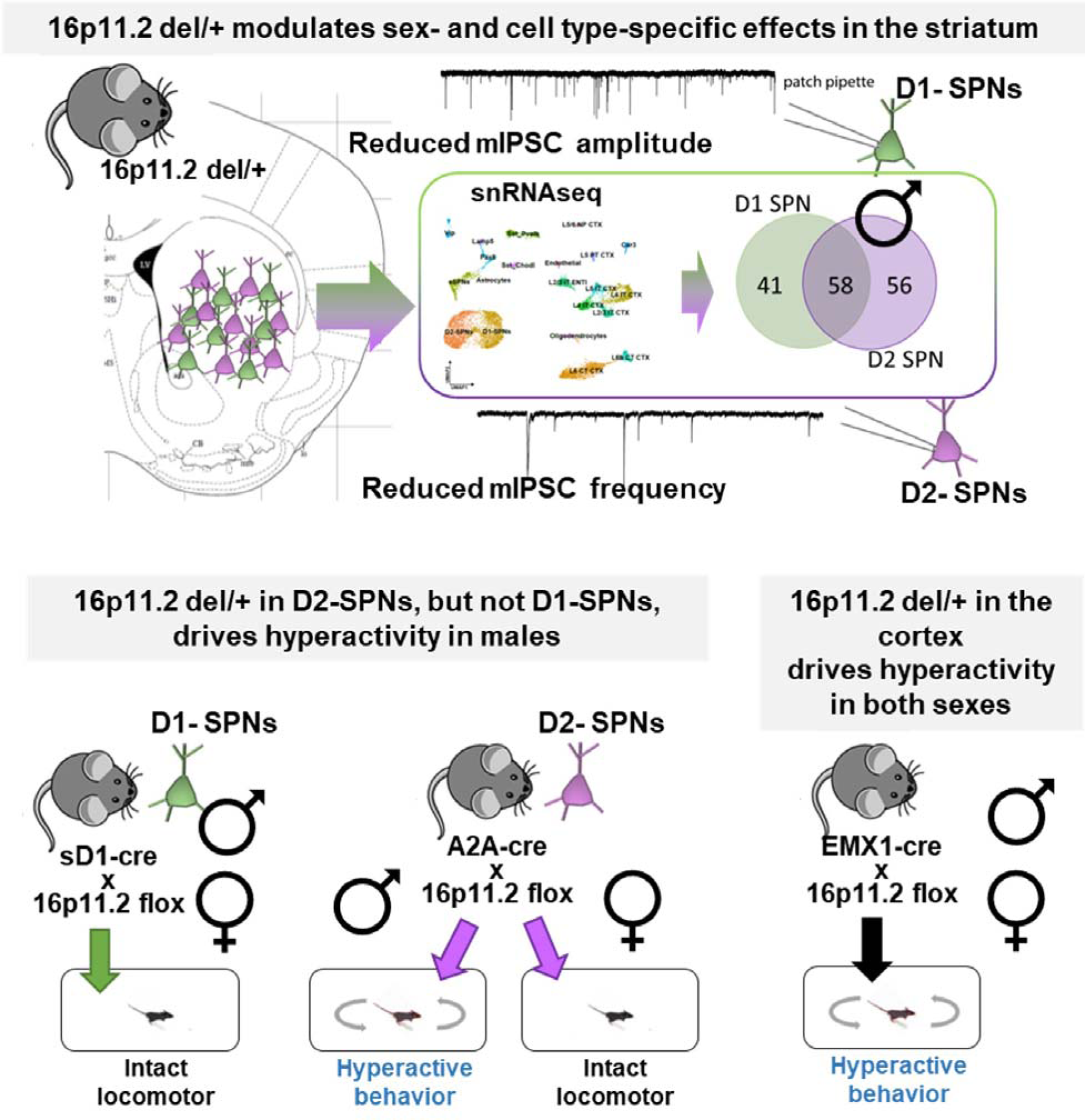

## Introduction

Male and female brains differ fundamentally in their reaction to genetic risk for brain disorders [1, 2]. Sex differences in incidence, symptom presentation, and outcome have been found across biologically heterogeneous disorders, ranging from affective disorders to neurodegenerative diseases and primary brain tumors [3–5]. Neurodevelopmental disorders (NDDs), including autism spectrum disorders (ASD) and attention-deficit hyperactivity disorder (ADHD), provide a particularly striking example of sex-specific vulnerability, with male-to-female ratios of 4:1 ASD and 2:1 for ADHD [6–9]. Although the primary features of ASD are consistent across sexes, females can present differently than males, exhibiting fewer or harder-to-detect behavioral alterations [10–13]. This bias is most likely driven by a differential vulnerability of distinct neuronal circuits to genetic risk factors [14, 15], but identifying the exact underlying mechanisms remains a major challenge in the field.

Multiple lines of evidence have highlighted striatal circuits as key mediators of vulnerability to NDDs [16–19]. Neuroimaging studies in humans diagnosed with NDDs and in genetic mouse models have identified changes in striatal structure, function, and connectivity [17, 18, 20–28]. The striatum is a major hub that shapes an array of behaviors, including the regulation of locomotor activity [16, 29]. Impairments in motor activity and stereotyped movements have been consistently observed in individuals diagnosed with NDDs [16, 30–32]. The proper acquisition and execution of the locomotor behaviors demand a precisely honed interplay of striatal networks. In the striatum, two distinct neuronal circuits, the direct and indirect pathways, play a critical role in locomotion. The direct pathway is responsible for the initiation of actions and requires the activity of dopamine 1 receptor expressing spiny projection neurons (D1-SPNs), whereas the inhibitory indirect pathway recruits striatal dopamine 2 receptor expressing (D2-) SPNs [33–38]. Previous studies have suggested an imbalance between the direct and the indirect pathway in NDDs [39–42]; however, a comprehensive understanding of the role of these pathways in mediating sex-specific effects on behavior is lacking.

Here, we explore mechanisms underlying the male preponderance of NDDs by mapping the sex-specific consequences of a copy number variation (CNV) linked to NDDs on the function of striatal circuits. We focus on 16p11.2 hemideletion (16p11.2 del/+), a rare genetic condition caused by a hemideletion within the chromosome 16p11.2 locus. This is one of the most common genetic risk factors for NDDs [43, 44]. Carriers of this CNV may receive a variety of diagnoses including ASD and ADHD [45–48]. Interestingly, female biological sex may be a protective factor for 16p11.2 del/+ carriers [49]. 16p11.2 del/+ model mice also show several sex-specific phenotypes [50–52], including behaviors dependent upon striatal circuits. Male mice demonstrate impairments in reward learning and motivation [53] and sleep [52], whereas hyperactivity is observed in both sexes [52]. Although previous studies have provided an initial indication of striatal circuit dysregulation in 16p11.2 del/+ mice [53, 54], the exact underlying mechanisms of these phenotypes are not fully understood.

In this study, we examine the sex-specific impact of 16p11.2 del/+ on striatal circuits. High-throughput single-nucleus RNA sequencing (snRNA-seq) of neuronal nuclei from the striatum reveals that 16p11.2 del/+ induces distinct transcriptomic signatures in D1-SPNs and D2-SPNs between males and females. Furthermore, we observe cell type-specific changes in inhibitory synapse function of SPNs in 16p11.2 del/+ male mice that support our transcriptomic findings. Crucially, selective deletion of the 16p11.2 region in D2-SPNs, induces hyperactivity in male but not in female mice indicating sexually dimorphic effects of the 16p11.2 region. Interestingly, 16p11.2 del/+ in the cortex is associated with hyperactivity in both sexes. Together, these results reveal that circuit level, sex-specific mechanisms mediate the behavioral alterations relevant to NDDs induced by 16p11.2 del/+.

## Results

### 16p11.2 del/+ drives sex- and cell type-specific transcriptomic changes in SPNs

To delineate the impact of 16p11.2 del/+ on specific neuronal circuits in the striatum, we first examine the transcriptomic landscape at the single cell level using snRNA-seq (figure 1A). In a previous study, single-cell gene expression was analyzed using multiplex single cell-qPCR, however, neonatal brains from 16p11.2 del/+ mice were studied, and sex information was not provided [54]. Here, we investigate cell type-specific transcriptomic changes in the striatum of both male and female 16p11.2 del/+ and wild-type (wt) adult mice. Based on the transcriptional signature, we identify sub-populations of striatal neurons, including direct pathway spiny projection neurons (D1-SPNs), and indirect pathway spiny projection neurons (D2-SPNs), as well as various cortical neurons (figure 1B, S1, S2). D1- and D2-SPNs exhibit marked transcriptional differences between 16p11.2 del/+ and wt mice that differ between the sexes. In D1- SPNs, we find 99 differentially expressed genes (DEGs) between 16p11.2 del/+ and wt males compared to 18 in females, whereas in D2-SPNs, we identify 114 DEGs in males compared to 20 in females (figure 1C-H, table S1-4). These results suggest a differential impact of hemideletion of gens in 16p11.2 region on male and female SPNs, with a ∼5-fold greater impact in males. In males, 58 DEGs are shared by both types of striatal neurons (figure 1I). Importantly, genes whose expression is altered by the 16p11.2 del/+ are enriched in genes associated with ASD. Using the SFARI gene database, we find that 21 DEGs in D1-SPNs of 16p11.2 del/+ male mice overlapped with ASD risk genes, and 21 DEGs in D2-SPNs overlapped with ASD risk genes, including 12 overlapping genes in both types of SPNs (figure 1I). Among these 30 genes, several, such as Adcy5, Camk2b, Herc1, Grin2b, Ube3a, Lrp1, and Elavl3, have been reported to be associated with motor function [55–61].

**Figure 1.**
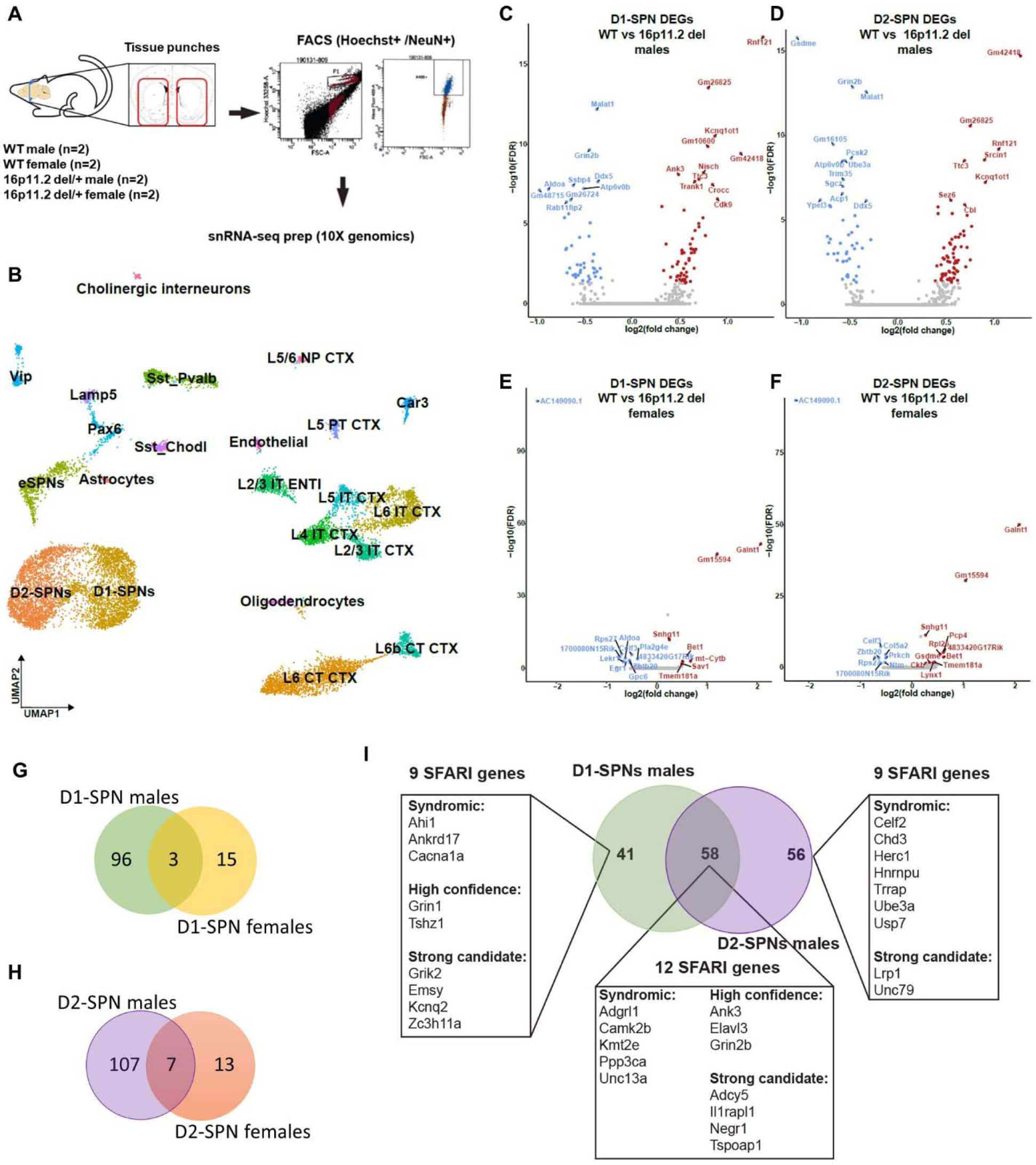
16p11.2 del/+ drives to sex- and cell-type specific transcriptomic changes in the striatum. (A) snRNA-seq preparation procedure. Tissue was harvested from male and female wt and 16p11.2 del/+ mice (n = 2). Dissection boundaries are indicated by red lines on the schematic illustrations. (B) Cell type annotation of the different cluster of neurons from the striatum and adjacent regions projected on a two-dimensional uniform approximation and projection (UMAP) map. The different clusters of cell types are labeled by their cell type identification name. (C, D) Differentially expressed genes in D-1 SPNs (C) and D2-SPNs (D) in 16p11.2 del/+ male mice compared to wt mice are displayed in volcano plots of statistical significance (in -log10) against fold-change (in log2) of gene expression change. Genes with a false discovery rate (FDR)□<□0.05 are highlighted in red (upregulated) and blue (downregulated) for significantly regulated. (E, F) Differentially expressed genes in D-1 SPNs (E) and D2-SPNs (F) in 16p11.2 del/+ female mice compared to wt mice are displayed in volcano plots. (G, H) Venn diagram representing the number of overlapping DEGs (adjusted p-value < 0.05 and log2FC > |0.25|) in D1-SPNs (C, E) or in D2-SPNs (D, F) of 16p11.2 del/+ mice relative to wt mice. (I) Venn diagram representing the overlap of significant DEGs (male D1- and D2-SPNs) from this study with the SFARI Gene autism susceptibility database focusing on the human gene module. The overlapping genes are represented in the pie charts according to their respective score developed by SFARI.

### 16p11.2 del/+ leads to distinct changes in synaptic properties in D1-SPNs and D2-SPNs in males

To characterize the molecular functions of the DEGs from D1- and D2-SPNs on male 16p11.2 mice, we perform pathway analysis. GO enrichment network analysis reveals that genes differently expressed in both D1- and D2-SPNs (58 DEGs) are associated with regulation of postsynaptic organization (figure 2A). These genes are more linked to the excitatory post-synapse organization. Consistently, in a previous study, increased miniature excitatory postsynaptic currents (mEPSCs) frequency and increased AMPAR/NMDAR ratio have been reported in 16p11.2 del/+ SPNs [54], suggesting both pre- and postsynaptic alterations at excitatory synapses. Interestingly, there is a substantial enrichment of genes associated with gamma-aminobutyric acid (GABA) synaptic transmission among the 41 exclusive D1-SPNs DEGs (figure 2A). Therefore, we investigate inhibitory synaptic function by recording spontaneous miniature inhibitory postsynaptic currents (mIPSCs) in 16p11.2 del/+ male mice. A comparison of mIPSC amplitudes reveals significantly reduced mIPSC amplitude in D1-SPNs of male 16p11.2 del/+ mice compared to wt mice (figure 2B, C) along with alterations in half-width and decay time of the synaptic currents (figure S3), suggesting a decrease in postsynaptic responsiveness of GABAergic synapses in D1-SPNs of 16p11.2 del/+ male mice (figure 2F). However, we do not detect changes in mIPSC amplitude onto D2-SPNs (figure 2D). Instead, comparison of mIPSC frequency reveals a significant decrease in mIPSC frequency in D2-SPNs of 16p11.2 del/+ male mice (figure 2E, S3), suggesting either reduced presynaptic release probability of GABA or fewer inhibitory inputs onto D2 SPNs (figure 2G). Together, these results suggest a differential impact of the high-risk polygenic factor 16p11.2 del/+ on molecular pathways and synaptic functions in the two major neuronal circuits in the striatum.

**Figure 2.**
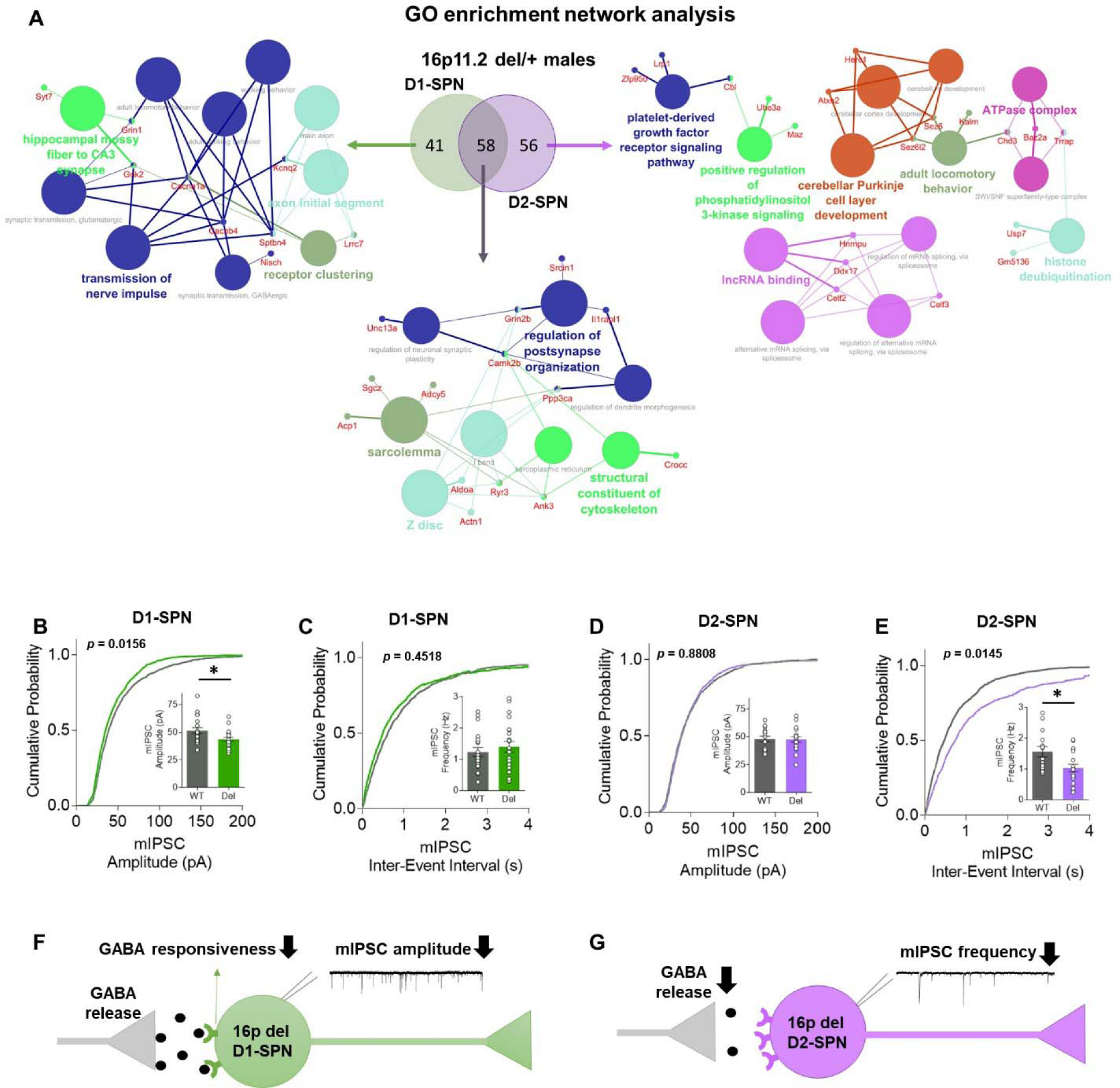
16p11.2 del/+ induces distinct changes in synaptic function in D1-SPNs and D2-SPNs in male mice. (A) Venn diagram representing the overlap of significant DEGs without the genes inside or near the 16p11.2 region between the D1-SPNs and D2-SPNs. Pathway analysis using ClueGo (KEGG and GO: Molecular function terms) of the male DEGs from either only D1-SPNs, only D2-SPNs or both D1-SPNs and D2-SPNs. Only pathways with a corrected p-value (Bonferroni step down) under 0.05 are displayed. The genes are labeled using CluePedia and are represented next to circles filled with either blue or red color to show the directionality of the dysregulation (blue = down-regulated; red = up-regulated). (B, C) Recording of synaptic events in NAc D1-SPNs of male 16p11.2 del/+ mice (n = 20 wt cells from 4 animals, n = 22 16p11.2 del/+ cells from 4 animals) show reduced mIPSC amplitude (p=0.0156, B), whereas the mIPSC frequency remained unchanged (p=0.4518, C). (D, E) Recording of synaptic events in NAc D2-SPNs of male 16p11.2 del/+ mice (n = 15 wt cells from 5 animals, n = 18 16p11.2 del/+ cells from 3 animals) show reduced mIPSC frequency (p=0.0145, E) with comparable mIPSC amplitude (p=0.8808, D). Error bars represent SEM. *p < 0.05. (F, G) Schematic diagrams of synaptic changes in D1-SPN (F) and D2-SPN (G) caused by 16p11.2 del/+ in males.

### Sex-specific transcriptomic changes in 16p11.2 del/+ male mice correlate with wildtype gene expression differences between sexes

To explore the effects of baseline sex differences on gene expression changes in male 16p11.2 del/+ mice, we compare gene expression between wt males and wt females to gene expression between wt males and 16p11.2 del/+ males (figure S4A, B). Interestingly, differentially expressed genes in 16p11.2 del/+ male mice positively correlate with the female gene expression profile in wildtype mice. We also apply the quadrant plot analysis to 16p11.2 del/+ females; however, the results are inconclusive because of the smaller number of DEGs (figure S4C, D). To investigate whether these gene expression patterns are specific to SPNs or observed more generally in the striatum, we analyze mRNA-seq data from a previous study (GSE224750) from the striatum of both male and female young adult 16p11.2 del/+ and wt mice (figure S4E, F). Quadrant plot analysis reveals similar results, indicating that sexual dimorphism at the molecular level between wt female and male mice correlates with male-specific gene expression changes induced by 16p11.2 del/+ (figure S4E). However, female gene expression changes induced by 16p11.2 del/+ are less affected by baseline differences between males and females (figure S4F).

### 16p11.2 del/+ in D2-SPNs leads to hyperactive behavior in male mice

To determine the impact of 16p11.2 del/+ on behavior linked to distinct striatal circuits, we utilize conditional genetic approaches to introduce the hemideletion selectively in either D1- or D2-SPNs. We generate mice with 16p11.2 del/+ exclusively in D2-SPNs by crossing 16p11.2 flox/+ mice and A2A-cre+/− mice (figure 3A). Double positive mice (flox/+:cre+/−; A2A-cre x 16p11.2 flox, D2-16p11.2 del/+ mice) are considered as the experimental group and the other three groups of mice (+/+:cre−/−, flox/+:cre−/−, +/+:cre+/−) are combined as controls. Young adult (3–4-month-old) male and female D2-16p11.2 del/+ mice and littermate control mice are monitored in the home-cage using an infrared beam-break system across the 24 hour cycle, similar to previous studies [52, 62]. We find increased activity in male (figure 3B, C, S5A) but not female (figure 3D, E, S5B) D2-16p11.2 del/+ mice, in contrast to our findings of hyperactivity in both sexes in the whole-organism 16p11.2 del/+ (figure S6) [52]. These results demonstrate that 16p11.2 del/+ in D2-SPNs induced hyperactive behavior specifically in male mice, revealing circuit level sex differences of the deletion of 16p11.2 region, consistent with our gene expression data.

**Figure 3.**
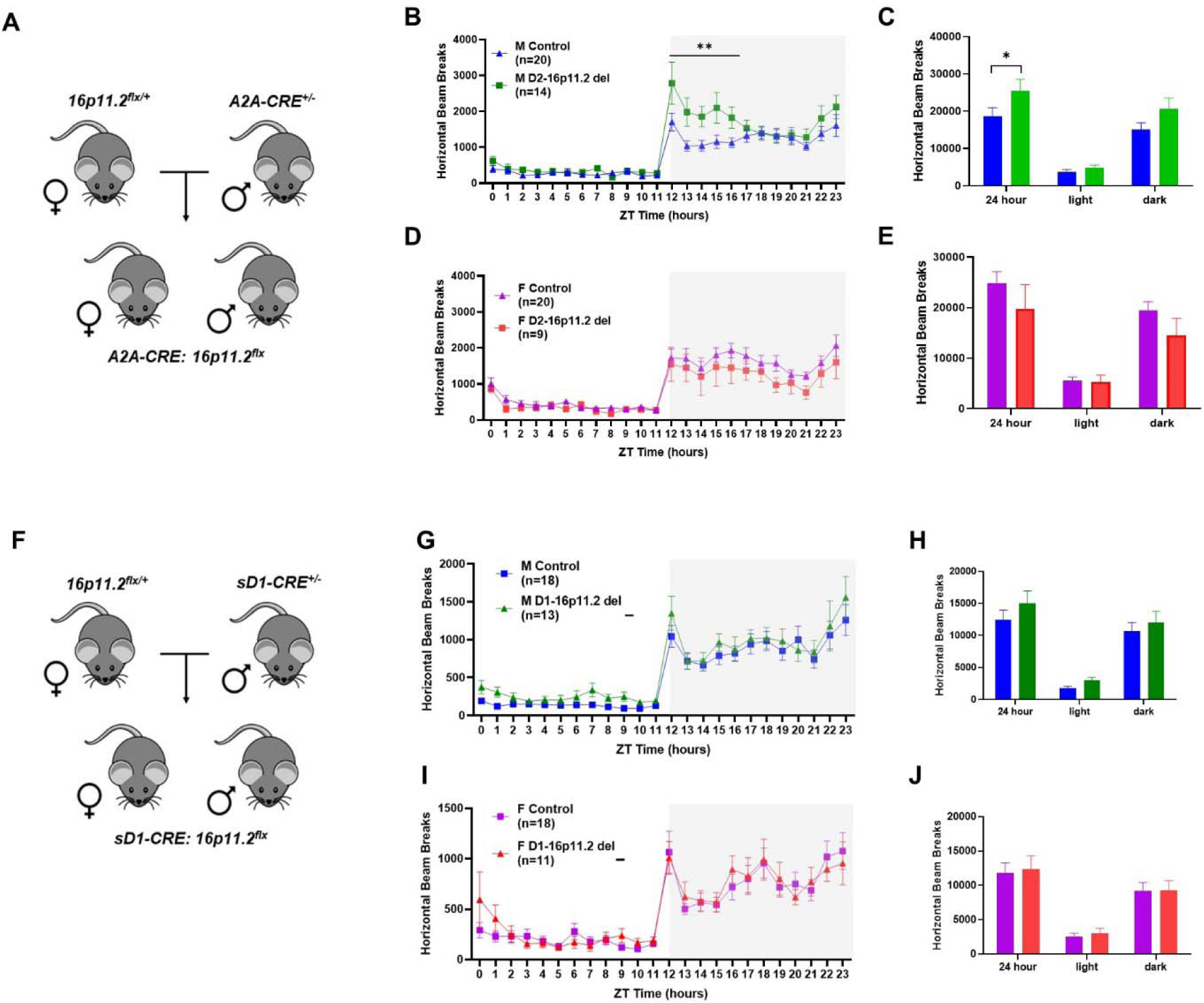
D2-SPN specific 16p11.2 del/+ mediates hyperactive behavior in males. (A) Genetic crosses used to induce 16p11.2 del/+ from D2-SPNs. (B-E) Activity monitoring shows increased activity in male A2A-cre x 16p11.2 flox mice. (B) Main effect of genotype, F(1, 32)=3.218, p=0.823; main effect of time, F(23, 736)=36.33, p<0.001; genotype x time interaction, F(23, 736)=2.694, p<0.001. Post hoc shows significant differences between male A2A-cre x 16p11.2 flox mice and control littermates in 12-16 hour time slots. (C) 24-hour plot shows increased activity in male A2A-cre x 16p11.2 flox mice (t=2.287, p=0.0244). (D) Main effect of genotype, F(1, 27)=1.263, p=0.2710; main effect of time, F(23, 621)=22.20, p<0.001; genotype x time interaction, F(23, 621)=0.5083, p=0.9738. (E) No change is reported in post hoc. (F) Genetic crosses used to induce 16p11.2 del/+ from D1-SPNs. (G-J) Activity monitoring results show equal locomotor activity regardless of genotypes in males (G, H) and females (I, J). Gray box indicates the dark cycle and mice activities are presented in 1-h bins (G, I). Mice activities are plotted by light/dark cycle (H, J). (G) Main effect of genotype, F (1, 29) = 1.137, p=0.2950; main effect of time, F(23, 667)=37.81, p<0.001; genotype x time interaction, F(23, 667)=0.4637, p=0.9856. (H) No change is detected in post hoc. (I) Main effect of genotype, F(1, 27)=0.06053, p=0.8075; main effect of time, F(23, 621)=25.27, p<0.001; genotype x time interaction, F(23, 621)=0.6693, p=0.8773. (J) No change is detected in post hoc. Error bars represent SEM. **p < 0.01.

To characterize the impact of 16p11.2 del/+ selectively in D1 SPNs, we generate mice with the hemideletion specifically in D1-SPNs by crossing 16p11.2 flox/+ mice and D1-cre+/− mice (figure 3F). Again, double positive mice (flox/+:cre+/−; sD1-cre x 16p11.2 flox, D1-16p11.2 del/+ mice) are considered as the experimental group and the other three littermate genotypes (+/+:cre−/−, flox/+:cre−/−, +/+:cre+/−) are categorized as the control group for behavioral tasks. Both male (figure 3G, H, S5C) and female (figure 3I, J, S5D) D1-16p11.2 del/+ mice exhibit similar locomotor activity to littermate controls. We confirm that neither 16p11.2 flox mice, D1-cre mice, nor A2A-cre mice exhibit hyperactive locomotion (figure S5, S7). Together, our results reveal that the sexually dimorphic effects of 16p11.2 del/+ on D2 SPNs mediates NDD-relevant increases in the locomotor activity of male animals.

Moreover, we find that risperidone, a potent antagonist of D2 receptors, alleviates the hyperactivity of 16p11.2 del/+ male mice (figure S8A). However, treatment with SCH39166, a D1 receptor antagonist, does not show similar effects (figure S8B), supporting the role of D2-SPNs in 16p11.2 del/+ male hyperactivity. These results support that 16p11.2 del/+ in D2-SPNs is a critical driver of hyperactive behavior in males.

### 16p11.2 del/+ in D2-SPNs in the dorsal lateral striatum drives hyperactive behavior in male mice

Distinct behavioral roles have been characterized for specific subregions within the striatum, with dorsal medial striatum (DMS) supporting more cognitive functions and dorsal lateral striatum (DLS) supporting motor output [16, 63]. However, the conditional genetic approaches using D1-cre and A2A-cre mice lack the specificity necessary to target neuronal circuits within specific subregions of the striatum. To target specific subregions within the striatum, we delivered SPN-subtype specific cre-virus to striatal subregions in 16p11.2 flox/+ male mice. AAV-ENK-cre virus contains the cre recombinase driven by the promoter for the enkephalin gene, which is specifically expressed in D2-SPNs [64, 65]. Injecting this virus in 16p11.2 flox mice localizes the 16p11.2 hemideletion effects to D2-SPNs within specific striatal subregions. By co-injecting a mix of AAV-ENK-cre and AAV-flex-tdTomato or AAV-flex-GFP in D2-GFP or D1-tdTomato reporter mice, we confirm AAV-ENK-cre virus expression exclusively in D2-SPNs (figure 4A-F). Then, we co-inject a mix of AAV-ENK-cre and AAV-flex-tdTomato in wt and 16p11.2 flox/+ male mice in three different striatal regions: DMS, DLS, and ventral striatum, and examine the impact on locomotor activity (figure 4G). Activity monitoring data shows that hyperactivity resulted from 16p11.2 del/+ in D2 SPNs in the DLS (figure 4I, J), but not in the DMS or ventral striatum (figure 4L, M, O, P). Following behavioral testing, brains are collected, and we confirm hemideletion of the 16p11.2 region in the striatum using PCR and verified injection location and the extent of viral expression using fluorescent microscopy (figure 4H, K, N, S10). In sum, these results demonstrate that 16p11.2 del/+ in D2-SPNs, specifically of the DLS, induces hyperactive behavior in male mice.

**Figure 4.**
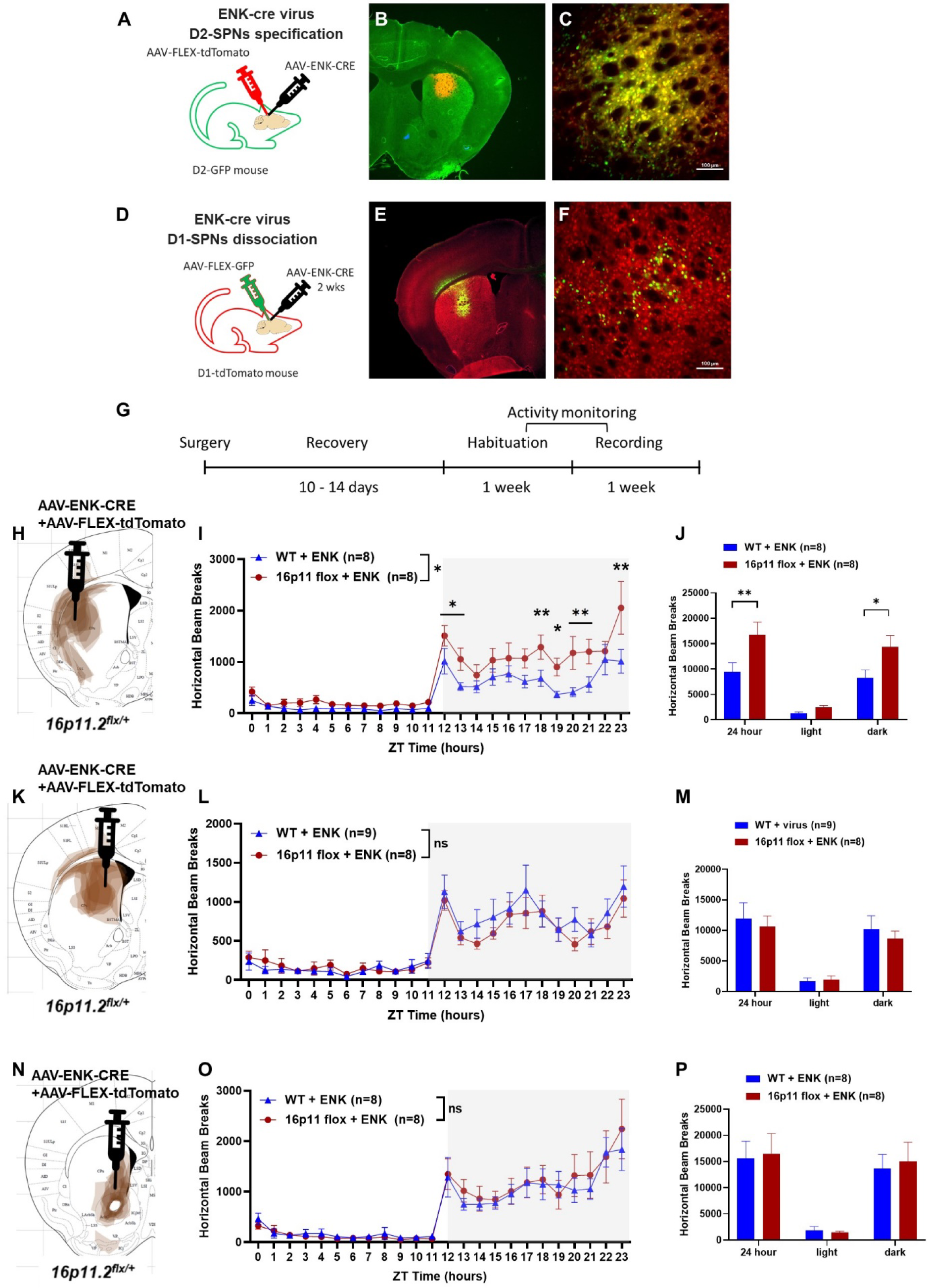
16p11.2 del/+ in D2-SPNs in the dorsal lateral striatum induces hyperactive behavior in males. (A) Schematic illustrates the virus mix injection of AAV-ENK-cre and AAV-FLEX-tdTomato into the striatum of D2-GFP mouse. (B) Representative fluorescence microscope images of AAV-ENK-cre and AAV-FLEX-tdTomato expression under 2.5x. (C) Representative fluorescence microscope images of AAV-ENK-cre and AAV-FLEX-tdTomato expression on D2-GFP mouse striatum (under 20x). ENK virus expression (red) is colocalized to D2-SPNs (green, ∼90% colocalization). Scale bar indicates 100um. (D) Schematic illustrates the virus mix injection of AAV-ENK-cre and AAV-FLEX-GFP into the striatum of D1-tdTomato mouse. (E) Representative fluorescence microscope images of AAV-ENK-cre and AAV-FLEX-GFP expression on D1-tdTomato mouse (under 2.5x). (F) Representative fluorescence microscope images of AAV-ENK-cre and AAV-FLEX-GFP expression on D1-tdTomato mouse striatum (under 20x). ENK virus expression (green) is dissociated to D1-SPNs (red, less than 15% colocalization). (G) Schematic of virus injection and behavioral task timelines. (H) Schematic drawing of virus expression in the dorsal lateral striatum across experiments in the current study (n=8). Red area represents the approximate virus expression. The position of the coronal section is 0.62□mm anterior to Bregma. (I) Main effect of genotype, F(1, 14)=5.577, p=0.033; main effect of time, F(23, 322)=21.07, p<0.001; genotype x time interaction, F(23, 322)=1.951, p<0.01. Post hoc shows significant differences between DLS AAV-ENK-cre injected 16p11.2 flox mice and control littermates in several time slots including 12-13, 18-21, and 23. (J) 24-hour plot and dark cycle plot show increased activity in DLS AAV-ENK-cre injected 16p11.2 flox mice (24-hour plot, t=3.032, p=0.0041; dark cycle plot, t=2.533, p=0.0151). (K) Schematic drawing of virus expression (red) in the dorsal medial striatum across experiments in the current study (n=7, 0.62□mm anterior to Bregma). (L) Main effect of genotype, F(1, 15)=0.1754, p=0.6813; main effect of time, F(23, 345)=24.20, p<0.001; genotype x time interaction, F(23, 345)=0.7224, p=0.8229. (M) No change is detected in post hoc. (N) Schematic drawing of virus expression (red) in the ventral striatum across experiments in the current study (n=7). The position of the section is 1.18□mm anterior to Bregma. (O) Main effect of genotype, F(1, 14)=0.0317, p=0.8613; main effect of time, F(23, 322)=23.55, p<0.001; genotype x time interaction, F(23, 322)=0.3469, p=0.9981. (P) No change is detected in post hoc. Error bars represent SEM. *p < 0.05, **p < 0.01.

### 16p11.2 del/+ in cortex causes hyperactivity in both male and female mice

16p11.2 del/+ in D2-SPNs drives hyperactive behavior only in male mice, suggesting sexual dimorphism on the circuit level. However, the circuit basis for the hyperactivity observed in whole-organism 16p11.2 del/+ female mice (figure S6, [52]) remains unresolved. We hypothesized that a loss of the 16p11.2 region in neurons within the cortex, one of the main excitatory inputs into the striatum, contributes to hyperactivity in female 16p11.2 del/+ mice. To explore the functional role of genes in the 16p11.2 region in the cortex, we crossed EMX1-cre mice and 16p11.2 flox/+ mice (figure 5A). As before, double positive mice (flox/+:cre+/−; EMX1-cre x 16p11.2 flox, Ctx-16p11.2 del/+ mice) are considered as the experimental group and the other three genotypes (+/+:cre−/−, flox/+:cre−/−, +/+:cre+/−) are combined as controls. Remarkably, both male (figure 5B, C) and female (figure 5D, E) Ctx-16p11.2 del/+ mice display hyperactive behavior compared to sex-matched control mice, phenocopying the 16p11.2 del/+ mice hyperactivity observed in both sexes (figure S6) [52]. These behavioral results demonstrate that deletion of the 16p11.2 region differentially impacts various hubs within cortico-striatal circuits, suggesting that circuit level sex differences mediate behavioral phenotypes of NDDs.

**Figure 5.**
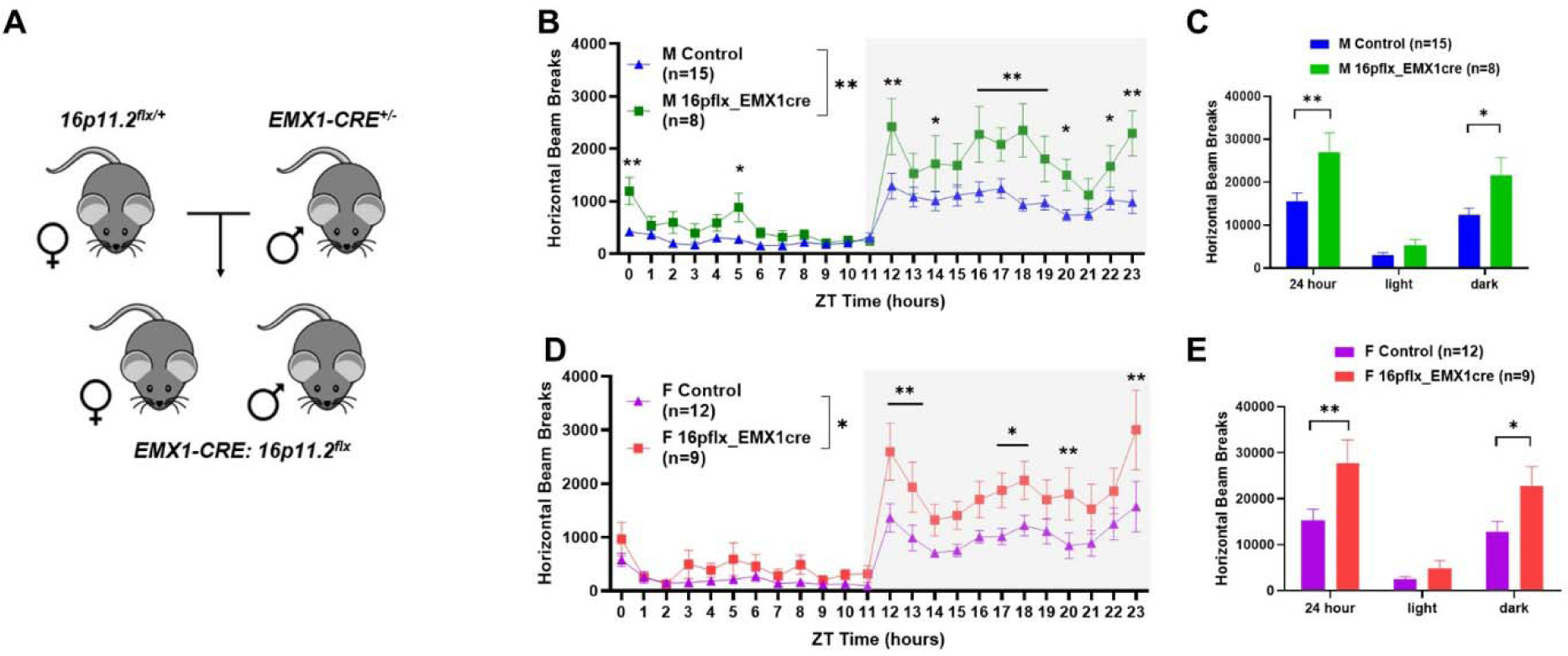
Cortex-specific 16p11.2 del/+ drives hyperactive behavior in both sexes. (A) Genetic crosses used to induce 16p11.2 del/+ in the cortex. (B, C) Activity monitoring shows increased activity in male EMX1-cre x 16p11.2 flox mice. (B) Main effect of genotype, F(1, 21)=9.199, p=0.0063; main effect of time, F(23, 483)=24.20, p<0.001; genotype x time interaction, F(23, 483)=2.984, p<0.001). Post hoc shows significant differences between male EMX1-cre x 16p11.2 flox mice and control littermates in several time slots. (C) 24-hour plot (t=3.524, p=0.0024) and dark plot (t=2.834, p=0.0185) show increased activity in male EMX1-cre x 16p11.2 flox mice. (D, E) Female EMX1-cre x 16p11.2 flox mice show hyperactivity compared to control littermates. (D) Main effect of genotype, F(1, 19)=5.815, p=0.0262; main effect of time, F(23, 437)=20.81, p<0.001; genotype x time interaction, F(23, 437)=1.785, p=0.0147). Post hoc shows hyperactivity of female EMX1-cre x 16p11.2 flox mice in several time slots. (E) 24-hour plot (t=3.083, p=0.0095) and dark plot (t=2.487, p=0.0475) show increased activity in female EMX1-cre x 16p11.2 flox mice.

## Discussion

In this study, we describe sex-specific contributions of the 16p11.2 del/+ region to both molecular and behavioral aspects of striatal circuits. Single-cell level transcriptomic analysis reveals substantial sex differences in the impact of 16p11.2 del/+ on D1- and D2-SPNs, as well as differences across other cell types. Conditional 16p11.2 del/+ in specific striatal circuits and in specific striatal subregions induces hyperactivity in a sex-specific manner, indicating sexually dimorphic effects of the genes in the 16p11.2 region on the function of distinct neuronal circuits within the striatum, underlying hyperactivity in 16p11.2 del/+ mice.

### Conditional 16p11.2 del/+ in distinct striatal pathways reveals circuit-level sex differences

Given the hyperactive behavior of 16p11.2 del/+ mice in both sexes, it is noteworthy that 16p11.2 del/+ in D2-SPNs drives different locomotor behavior phenotypes between male and female mice. These results indicate that 16p11.2 del/+ in D2-SPNs is a critical driver of hyperactive behavior in males, but not in females, revealing circuit-level sex differences underlying hyperactivity in 16p11.2 del/+ mice. This is significant because it implies that even when male and female mice exhibit the same behavioral phenotype, different neuronal circuits may underlie this phenotype in males and females. There has been substantial interest in profiling sex differences in striatal anatomy and function. Clinical reports have shown sex differences in dopamine release at baseline in the ventral striatum [66]. In rodent models, it has been reported that dopamine release and uptake dynamics, as well as the expression of dopamine receptors in the striatum, differ depending on sex [67–70]. Additionally, several preclinical studies have shown sex-dependent responses to psychostimulants in the striatum [67, 71, 72]. Sex differences in the distribution and density of striatal interneurons are reported [73]. Together, our findings of circuit-level sexually dimorphic effects of the genes in the 16p11.2 region suggest that different therapeutic approaches for females and males, even when they exhibit the same behavioral phenotype and carry the same genetic deletion. Further support for sex-specific circuit effects that define behavior is provided by our finding that 16p11.2 del/+ in cortical excitatory neurons phenocopy whole-organism 16p11.2 del/+ mice hyperactivity observed in both sexes. These findings indicate that deletion of the 16p11.2 region has a differential impact on key hubs within cortico-striatal circuits in a sex-specific manner. The striatum receives a complex array of sensory and contextual information from cortical afferents, and the behavioral outputs are subject to the input sent from cortical regions that the striatum receives [63, 74, 75]. Moreover, a recent study showed that disruption of the hyper-direct pathway, a circuit of cortico-subthalamic projections bypassing the striatum, induces hyperactive behavior in mice [76]. Therefore, investigating the cortex-specific mechanisms underlying male and female hyperactivity will provide further insight into the circuit-level sex differences mediating hyperactive behavior in 16p11.2 del/+ mice.

### D2-SPNs is a critical driver of hyperactive behavior in 16p11.2 del/+ males

The traditional rate model of movement control suggests that the two major neuronal circuits in the striatum antagonize each other: the direct pathway (D1-SPNs) initiates movement, while the indirect pathway (D2-SPNs) suppresses it [35, 36, 77, 78]. However, recent studies have challenged this view, providing evidence that both pathways are activated during the initiation of movements, suggesting that the coordinated activity of both pathways is necessary for the control of movement [79–82]. Our results show that 16p11.2 del/+ in D2-SPNs, but not in D1-SPNs, induces hyperactive behavior in male mice, demonstrating the important role of D2-SPNs in 16p11.2 del/+ male hyperactivity. Additionally, we discover that risperidone, a potent antagonist of D2 receptors, and off-label prescription for the treatment of ADHD [83, 84], effectively reduces the hyperactivity observed in male 16p11.2 del/+ mice. Together, these findings underscore the pivotal role of 16p11.2 del/+ in D2-SPNs as a key contributor to hyperactivity in males.

### Conditional 16p11.2 del/+ in DLS, but not DMS, induces hyperactivity in males

Distinct roles have been proposed for the DLS and the DMS: the DLS is linked to habitual actions, while the DMS is associated with goal-directed behavior [63]. These distinct roles are associated with unique striatal afferent connections: Somatic sensorimotor subnetworks extend their projections into the DLS, while associative cortical areas, such as anterior cingulate cortex and prelimbic cortex, project into the DMS [16, 63, 75, 85]. In line with the distinct roles of the DLS and the DMS, our study reveals that 16p11.2 del/+ within D2-SPNs of the DLS, but not the DMS, results in hyperactive behavior. While we demonstrate the impact of genetic variations linked to NDDs on these subregions regarding behavioral phenotypes, the molecular impact of these genetic variations on striatal subregions, specifically the DMS and DLS, remain unexamined. To address this gap, future experiments, such as spatial transcriptomics, are essential for a comprehensive characterization of region-specific phenotypes.

### snRNA-seq analysis emphasizes the significance of polygenic influences of ASD risk genes

Recent research underscores the notion that transcriptomic patterns of brain disease risk genes offer a unique molecular signature specific to each disorder [86]. Additionally, it has been indicated that multiple genes within the 16p11.2 region may exert significant polygenic influences, not only in ASD but potentially across a broader spectrum of NDDs, solidifying the 16p11.2 locus as a wellspring of both common and rare genetic variations [87], while previous investigations have failed to pinpoint any single gene within the 16p11.2 locus strongly associated with ASD [88]. Our study delves into the intricate landscape of gene expression in this mutation cell-type specifically. Through single-cell level transcriptomic analysis, we unveil several DEGs within D1- and D2-SPNs associated with ASD based on the SFARI gene database, suggesting a potential compensatory mechanism in response to the mutation. The acquisition of additional genetic hits in the ASD risk genes could predispose carriers to illness, potentially elucidating the phenomenon of healthy carriers. This mechanism also provides insights into the pleiotropy observed in CNVs, where carriers exhibit an increased nonspecific risk. In essence, while carriers of CNVs may initially appear healthy due to compensatory mechanisms, the accumulation of additional mutations heightens their susceptibility to diverse disorders, ultimately leading to clinical manifestations. In this context, our snRNA-seq analysis contributes to the growing body of literature elucidating the polygenic effects of ASD risk genes, providing a comprehensive framework for unraveling the complex genetic underpinnings of NDDs.

### snRNA-seq analysis identifies genetic factors associated with hyperactive behavior of 16p11.2 del/+ male mice

In our study, single-cell level transcriptomic analysis reveals a substantially different impact of 16p11.2 del/+ on D1- and D2-SPNs. Interestingly, pathway analysis of snRNA-seq reveals that DEGs in both D1- and D2-SPNs are associated with adult locomotory behavior. However, only 16p11.2 del/+ in D2-SPNs mediates hyperactive behaviors. In D2-SPNs, we identify several DEGs that are enriched in the adult locomotory behavior-related pathways and other genes associated with motor regulation or movement deficits. Specifically, *Pde1b*, highly expressed in the striatum and involved in baseline motor activity [89], is upregulated in D2-SPNs in 16p11.2 del/+ mice, indicating a potential molecular mechanism driving hyperactivity. Furthermore, *Atxn2* and *Mtcl2* are associated with motor deficits [90–92]. In addition, comparisons between ASD risk genes and DEGs in D1- or D2-SPNs reveal that 30 DEGs are enriched in 16p11.2 del/+ male mice. Of these, 7 genes (*Adcy5, Camk2b, Herc1, Grin2b, Ube3a, Lrp1*, and *Elavl3*) have been reported to be associated with motor function [55–61]. All seven of these genes exhibit enrichment in D2-SPNs, with four showing enrichment in both D1- and D2-SPNs. Genes identified in the adult locomotory behavior-related pathway in D2-SPNs, as indicated by GO enrichment network analysis, along with genes enriched in the SFARI gene database, underscore the multifaceted influence of these genetic factors on motor function. Interestingly, this broad effect of 16p11.2 deletion on the expression of a significant number of autism-associated genes potentially serves as an explanation for the polygenic nature of autism and suggests that the study of single locus mutations might have a polygenic impact. Investigating the circuit-specific impact of these genes in the striatum will be crucial for future studies, enhancing our understanding of potential mechanisms for hyperactive locomotion.

### Molecular level sexual dimorphism in striatal circuits

Our snRNA-seq analysis reveals sex-specific impacts of 16p11.2 del/+ in D1- and D2-SPNs, demonstrating a substantial number of DEGs in males compared to females. Interestingly, our results indicate that molecular level sexual dimorphism in the striatum between wt female and male mice correlates with male-specific gene expression changes induced by 16p11.2 del/+, displaying an unexpected shift of the gene expression patterns in 16p11 del/+ males towards wt females. Physiologically, striatal networks differ between sexes, as evidenced by sexual dimorphism in the development of striatal circuits in healthy youth, particularly within cortico-striatal-thalamic circuits [93–95]. Furthermore, rodent studies have reported sexual dimorphism in neurotransmission, neuronal excitability, and quantity of neuronal subtypes participating in these circuits [96–99]. Additionally, RiboTag RNAseq studies have highlighted molecular-level sex differences in the striatum, demonstrating moderate sexual dimorphism in D1- and D2-SPNs at baseline [100]. Despite several studies demonstrating sexual dimorphism in NDDs, the understanding of how baseline sexual dimorphism interacts with genetic variation linked to NDDs, leading to sex-specific behavioral phenotypes, is still incomplete. We believe our findings provide a new perspective on the factors influencing male vulnerability or female resilience to NDDs by connecting molecular sexual dimorphism to sex-specific transcriptomic changes and may alert others to consider baseline sex differences more closely in their research.

### Impairments in striatal inhibitory synaptic transmission in 16p11.2 del/+ males

In alignment with our observed cell type-specific transcriptomic changes, acute slice electrophysiology experiments reveal a cell type-specific impairment in inhibitory synaptic transmission in 16p11.2 del/+ mice. We detected reduced amplitude of inhibitory synaptic postsynaptic currents onto D1-SPNs, along with alterations in half-width and decay time of the synaptic currents. Changes in mIPSC amplitude could be attributed to fewer postsynaptic receptors or changes in their subunits or phosphorylation [101]. These results are consistent with postsynaptic changes, specifically at the receptor level. In contrast, we found decreased frequency of inhibitory synaptic currents onto D2-SPNs, suggesting a presynaptic change, although these results do not address the source of inhibitory input information for D2-SPNs. Moreover, a previous study noted increased sEPSC frequency in D2-SPNs in 16p11.2 del/+ mice [54]. Together these results suggest an overall increased output from D2-SPNs. In accordance with the cell type-specific transcriptomic alterations, our electrophysiology experiments support cell type-specific changes in synaptic properties of 16p11.2 del/+ mice.

### Overall conclusions

This study demonstrates that a CNV associated with NDDs acts in distinct ways in different neuronal circuits, impacting gene expression, synaptic function, and behavior in a sex-specific fashion. These findings provide new insight into male vulnerability or female resilience to NDDs, highlighting striatal circuits as key mediators of symptoms of NDDs and provide new avenues for circuit-based, sex-specific therapeutic approaches.

## Materials and Methods

### Animals

The 593 kb CNV (chr6, 133.84-134.28 Mb) conditioned mouse line was generated in the Mills laboratory to delete the causative region in specific neurons of the developing brain using chromosome engineering [102, 103]. This conditional model has a floxed version of the region analogous to 16p11.2, and when crossed to a cell-type specific cre mouse model, causes efficient deletion of the floxed region in vivo [104, 105]. 16p11.2 del/+ mice were produced by breeding male B6129S-Del(7Slx1b-Sept1)4Aam/J mice purchased from The Jackson Laboratory (Stock #013128) with female B6129SF1/J mice (Stock #101043). For cell-type specific electrophysiology approach, we crossed 16p11.2 del/+ male mice with Drd1a-tdTomato BAC transgenic (D1-tdTomato) female mice (purchased from GENSAT, Tg(Drd2-EGFP)S118Gsat/Mmnc, Stock Number 000230-UNC) or Drd2-EGFP BAC Transgenic (D2-GFP) female mice (purchased from The Jackson Laboratory, B6.Cg-Tg(Drd1a-tdTomato)6Calak/J, Stock Number 016204). To generate mice with cell type-specific deletion of the region analogous to 16p11.2 (the 7F4 region in mice) specifically in D1- or D2-SPNs, we used the EY217 Drd1a-cre line (purchased from GENSAT, B6.FVB(Cg)-Tg(Drd1a-re)EY217Gsat/Mmucd, Stock Number :034258-UCD) and KG139 A2A-cre line (purchased from GENSAT, B6.FVB(Cg)-Tg(Adora2a-cre)KG139Gsat/Mmucd, Stock Number 036158-UCD). These cre lines were backcrossed at least 5 generations to 129S1 (purchased from The Jackson Laboratory, 129S1/SvImJ, Stock Number 002448) and bred to floxed 16p11.2 hemizygous mice created by our collaborator Alea Mills at Cold Spring Harbor Laboratory [103]. To selectively introduce the 16p11.2 del/+ in the cortex, we used Emx1-cre line (purchased from The Jackson Laboratory, B6.129S2-Emx1tm1(cre)Krj/J, Stock Number 005628). Emx1-cre mice express the cre recombinase in cortical excitatory neurons and glia, but not GABAergic neurons [106]. To selectively introduce the 16p11.2 del/+ in D2-SPNs, we utilized A2a-cre mice instead of D2-cre mouse. It is because cre recombinase is expressed in striatal cholinergic interneurons as well as D2-SPNs [107], unlike A2a-cre mice, which directs D2-SPNs but not cholinergic interneurons [36]. Experimental data was collected from littermate animals during the same time period. Mice aged between 10 and 16 weeks were randomly used for experiments to minimize any bias unless otherwise specified.

Mice were housed in a room with a 12-hour light/dark cycle and were allowed *ad libitum* access to food and water. All procedures were approved by the University of Iowa Institutional Animal Care and Use Committee, and followed policies set forth by the National Institutes of Health Guide for the Care and Use of Laboratory Animals.

#### Genotyping

PCR-genotyping was done to identify founders with the intended mutation.

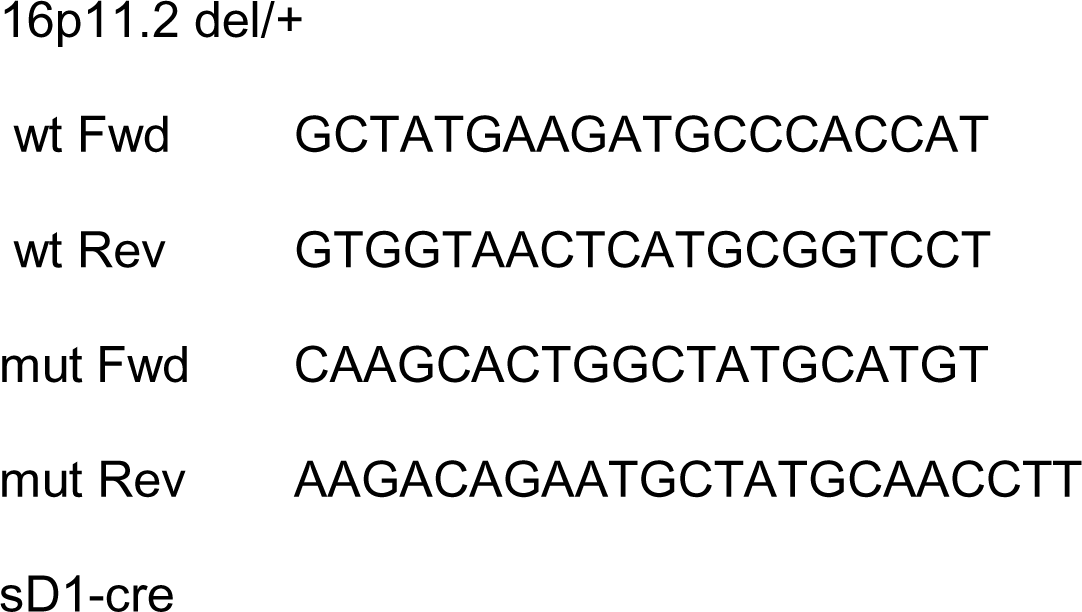
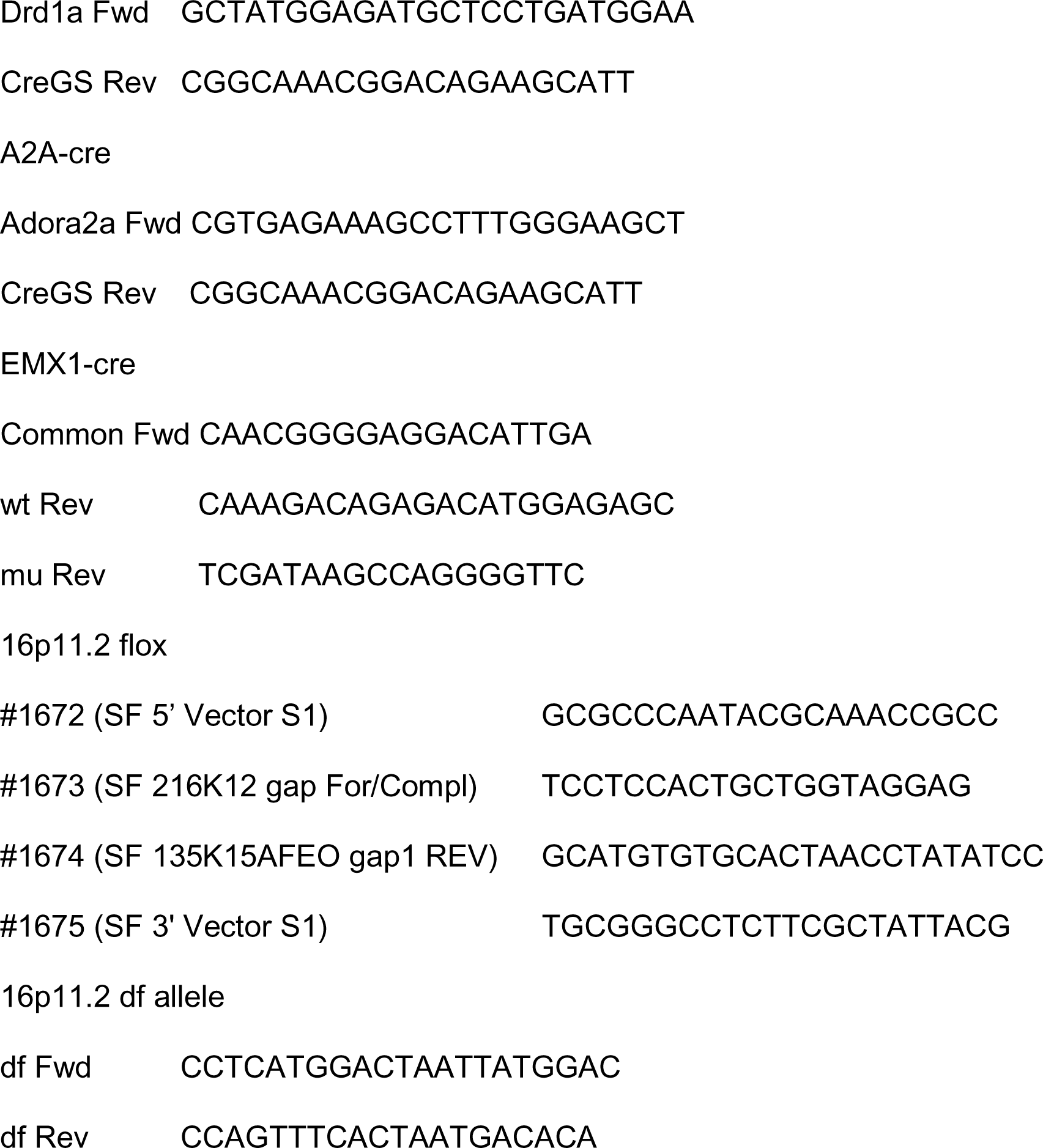

### Single-cell RNA-sequencing

#### Tissue processing for scRNA-seq

Mice were sacrificed at 14-16 weeks of age by live decapitation and the brain was rapidly removed. Using a mouse brain matrix, 1 mm coronal sections were collected, and then tissue punches containing the striatum (−0.5 ∼ 1.5 mm AP from bregma; n=2 mice/genotype/sex) were rapidly frozen on dry ice. Tissue punches were preserved at −80°C until the day of single nuclei dissociation.

#### Single nuclei purification

For single nuclei dissociation, a protocol from previous study was followed [108]. Briefly, frozen tissue punches were chopped and transferred to 450ul of chilled (4°C) homogenization buffer (250mM sucrose, 25mM KCL, 5mM MgCl2, 10mM Tris buffer (pH 8.0), 1μM DTT, 1x protease inhibitor, RNaseIn 0.4 U/ul, Superasin 0.2 U/ul, 0.1% Triton X-100). After the Dounce homogenization and filtering (40μm), the nuclei were concentrated by centrifugation (1,000g for 8 min at 4°C). For staining, anti-NeuN antibody, Hoeshst, and Alexa 488 secondary antibody were treated and incubated on rocker in cold room for 30 min. The nuclei were resuspended in 600 ul FACS buffer (1% unacetylated BSA, RNaseIn 0.4 U/ul, PBS) for FACS to further purify the nuclei for sequencing. The nuclei were first sorted based on Hoechst signal and then based on Alexa 488 fluorophore, collecting 50,000 nuclei per mouse.

#### Single Nuclear (snRNA) RNA Sequencing and Analysis

Libraries were constructed using the 3’ Expression-Single-Cell RNA-Seq using 10X Chromium System (v3.1), according to the manufacturer’s instructions. The nuclei were brought to a concentration of 500 nuclei/μl and 5,000 nuclei were loaded to prepare libraries. Libraries were then sequenced (pair-end sequencing) on the Illumina NovaSeq6000 at the Iowa Institute of Human Genetics at University of Iowa. The depth of sequencing per flow cell was 50,000 reads per nuclei. FASTQ files were aligned to the pre-mRNA annotated Mus musculus reference genome version GRCm38 (refdata-cellranger-mm10-3.0.0). Cell Ranger Count v3.0.2 (10x Genomics, Pleasanton CA) was used on FASTQ data from each of the GEM wells individually for genome alignment and feature-barcode matrix counts generation. Cell Ranger Aggr v3.0.2 was then used to combine data from multiple samples of the same genotype (biological replicates) into an experiment-wide feature-barcode matrix and analysis and normalizing those runs to the same sequencing depth and then recomputing the feature-barcode matrices and analysis on the combined data. This raw expression UMI counts matrix contains cells as rows and genes as columns and can be further used for downstream analysis such as normalization, clustering, differentially expressed genes, etc.

The clusters were then visualized with uniform manifold approximation and projection (UMAP) in two dimensions of Unique Molecular Identifiers (UMI) per cells (figure S1B, C). The most significant gene markers corresponding to each cluster and conserved across the two genotypes (WT and 16p11.2 del/+) were used to identify the cell-types by correlating with previously published single-cell data [109] and from the transcriptomic explorer of the Allen Brain Atlas (https://celltypes.brain-map.org/rnaseq/mouse_ctx-hip_10x). To obtain an observer-independent data-driven characterization of cell types, we computed the 2000 most variable genes in each cell, based on the assumption that these genes will distinguish best between different cell type and formed clusters of specific cell types based on similar transcriptomic signatures for each cell (figure 1B, S1, S2). DotPlot representations of the normalized UMI expression of conserved gene markers across the two genotypes in the different cell types.

Raw single nuclei RNA-seq UMI count data was used for clustering analysis using Seurat R analysis pipeline [110]. First, cells with more than 50,000 molecules (nUMI per cell) were filtered out to discard potential doublets cells. Post filtering, the raw UMI counts from primary filtered dataset were used for log-normalization and scaled using a factor of 10,000 and regressed to covariates such as number as described in Seurat analysis pipeline [110]. To further identify the top variable genes, the data was used to calculate principal components (PCs). Using Jackstraw analysis, statistically significant PCs were used to identify clusters within the data using original Louvain algorithm as described in Seurat analysis pipeline followed by visualizing the clusters with uniform manifold approximation and projection (UMAP) in two dimensions of UMI per cells [111]. Genes corresponding to each cluster were used to identify the cell-type by correlating to genes expressed in previously published adult mouse striatal single cell data [109] and analyzed for expression of top marker genes of known cell-types. Pairwise differential gene expression analysis tests were performed within each cluster-pair (wild-type VS 16p11.2 del/+) using a Wilcoxon Rank Sum test from the Seurat R analysis pipeline [110] to identify genes differentially expressed. A significant threshold was setup for genes with an adjusted p-value under 0.05 and an absolute log_2_ fold change above 0.2. Enrichment analysis of DEG-associated pathways and molecular functions from snRNA-seq was performed with a combination of Kyoto Encyclopedia of Genes and Genomes (KEGG) and the Gene Ontology (GO-Molecular Function-EBI-Uniport-GOA-) databases. The analyses were done with the Cytoscape (version.3.8.0, https://cytoscape.org/) [112] plug-in ClueGO (version 2.5.6) [113]. Only the pathways with a p-value < 0.05 and gene counts ≥ 3 were considered as significant and displayed. To connect the terms in the network, ClueGO utilizes kappa statistics in which here was set as ≥ 0.4.

### Electrophysiology

All analyses were performed on male littermate mice of 2-5 months of age. Mice were anesthetized with isoflurane and perfused transcardially with ice-cold artificial cerebrospinal fluid (ACSF) (pH 7.3-7.4) containing (in mM): 124 NaCl, 2.5 KCl, 1.2 NaH2PO4, 24 NaHCO3, 5 HEPES, 12.5 glucose, 1.3 MgSO4, 7H2O, 2.5 CaCl2. Subsequently, mice were decapitated, and brains were quickly removed. The brain was blocked off, removing a small portion of the left lateral hemisphere to create a flat surface for placement on the vibratome stage. Parasagittal slices (250µm) containing the nucleus accumbens (NAc) core were prepared on a vibratome (VT1200s, Leica). Slices were incubated in a holding chamber for 12-15 minutes at 32-34°C in a NMDG-based recovery solution (pH 7.3-7.4, pH adjusted with HCl) (in mM): 92 NMDG, 2.5 KCl, 1.2 NaH2PO4, 30 NaHCO3, 20 HEPES, 25 glucose, 5 sodium ascorbate, 2 thiourea, 3 sodium pyruvate, 10 MgSO4, 7H2O, 0.5 CaCl2. Osmolarity for the NMDG-based solution and ACSF was kept between 300-310 mOsm. Following incubation, slices were moved to a second holding chamber containing ACSF at room temperature (20-22°C) for at least 60 minutes prior to recording experiments. Slices were removed from the holding chamber and placed in the recording chamber (Scientifica) fully submerged in (95% O2, 5% CO2) ACSF at a perfusion rate of 1.4-1.6 mL/min, bath temperature of 29-30°C, and secured using a slice anchor (Warner Instruments).

#### Intracellular Recordings

Whole-cell voltage-clamp recordings from SPNs were obtained using IR-DIC video microscopy on an upright microscope (Olympus, BX51). The NAc core was identified by the presence of the anterior commissure as previously described [114, 115]. Targeted recordings of D1-tdTomato and D2-EGFP positive SPNs, were identified by the presence of the fluorophore that was excited with illumination from a collimated light-emitting diode (LED) illuminator (CoolLED, PE-300) through a 40x objective (Olympus, 0.8NA water immersion). Recordings were made with borosilicate glass (World Precision Instruments, TW150-3) with a tip resistance of 3-5MΩ filled with high internal chloride to maximize inhibitory current amplitude (in mM): 135 CsCl, 10 HEPES, 0.6 EGTA, 2.5 MgCl, 10 Na-Phosphocreatine, 4 Na-ATP, 0.3 Na-GTP, 0.1 spermine, 1 QX-314. All miniature synaptic currents were recorded in the presence of tetrodotoxin (500 nM). To isolate inhibitory currents, D-APV (50 µM) and NBQX (10 µM) were added to block NMDARs and AMPARs, respectively. Voltage-clamp recordings were done holding the cell at a membrane potential of −70mV using a MultiClamp 700B patch-clamp amplifier (Molecular Devices) filtered at 2.8kHz and digitized at 20kHz. Input resistance and access resistance were noted subsequently following membrane break-in, and cells with RA > 25 MΩ or RI > 300 MΩ were discarded from further analysis.

Data acquisition was performed using custom-built Recording Artist software (Dr. Rick Gerkin), Igor Pro 6.37 (Wavemetrics). Detection of miniature events used for analyses was performed in Mini-analysis (Synaptosoft). Data visualization and statistics was performed with GraphPad Prism v7.0. All data are presented as the mean ± SEM, with N referring to the number of animals and n to the number of cells.

### Activity monitoring

An infrared beam-break system (Opto M3, Columbus Instruments, Columbus, OH) was used to measure mice locomotor activity, as previously described [52]. Behavior experiments were designed based on previous experience with an effect size of 0.7 in a power analysis. The mice were acclimatized to the monitoring chambers for one week before data collection to reduce the effect of a new environment. The data was collected for one week following the habituation period. ANOVAs followed by Fisher’s LSD post-hoc tests were performed using GraphPad 9 (La Jolla, CA), with genotype as the between-subjects factor and time as the within-subjects factor, as previously described [52]. After behavioral assessments, the striatum and adjacent cortices were dissected from D1- and D2-16p11.2 del/+ and control mice to confirm the hemideletion of the 16p11.2 region in the striatum using PCR (figure S9).

### Virus Injection and stereotaxic injections

AAV-ENK-cre virus was kindly provided by Dr. Jocelyne Caboche, Sorbonne University. AAV-ENK-cre virus drives a D2-SPNs-specific expression of the cre recombinase [64, 65]. pAAV-FLEX-tdTomato and pAAV-FLEX-GFP viruses were used to confirm AAV-ENK-cre virus expression (purchased from Addgene, Catalog # 28306 and #28304, respectively). 16p11.2 flox mice were injected with the virus mix (10^12^ vg/animal for AAV-ENK-cre and 1×10^11^ vg/animal for either pAAV-FLEX-tdTomato and pAAV-FLEX-GFP) into the dorsal medial striatum (AP, +0.65□mm; ML, ±□1.70□mm; DV, −3.2□mm from bregma), dorsal lateral striatum (AP, +0.65□mm; ML,□±□2.2□mm; DV, −3.2□mm from bregma) or NAc (AP, +1.2 mm; ML, ±1.0 mm; DV, −4.2 mm from bregma). The rate of injection was 200 nl/min with a total volume of 1 μl per striatum. Following 2 weeks of virus expression, mice were tested in activity monitoring. After behavioral assessments, the striatum and cerebellum were dissected from 16p11.2 flox and control mice to confirm the hemideletion of the 16p11.2 region using PCR (figure S10).

### Fluorescence Imaging

After activity monitoring, mice were cervically dislocated and decapitated, and the whole brain removed on ice. Brain tissue was immersed in the 4% formaldehyde (PFA) at 4°C overnight. The tissue was transferred into 30% sucrose for 48 h. Then, tissue was sliced into 40 µm thick sections using a cryostat at −20 °C and mounted on slides. Fluorescence images were taken with a Leica DM6B Upright digital microscope using the software program NeuroLucida (Microbrightfield Inc).

## Data and code availability

The datasets generated and/or analyzed during the current study are available in the NCBI’s Gene Expression Omnibus repository, GEO Series accession GSE266222. The code for analyses and figures related to snRNA-seq data can be accessed through <u>GitHub - YannVRB/snRNA-seq-16p11.2-del-males-females</u>.

## Publication acknowledgement

This work was supported by The University of Iowa Hawkeye Intellectual and Developmental Disabilities Research Center (HAWK-IDDRC) P50 HD103556 (T.A. and Lane Strathearn, PI), the Roy J. Carver Chair in Neuroscience (T.A.), Interdisciplinary Graduate Program in Genetics at University of Iowa (Y.V.), NIH grant R01 MH 087463 (T.A.), NIH grant R01 DA 056113 (T.A.), Simons Foundation Autism Research Initiative (SFARI) grant 345034 (T.A.), NIH grant T32 GM067795 (B.K.), NIH grant F31 MH134542 (B.K.), NIH grant R01 MH115030 (M.F.), Eagles Autism Challenge (M.F. & T.N.-J.) and the Andrew H. Woods Professorship (T.N-J.).

The Neural Circuits and Behavior Core in the Iowa Neuroscience Institute provided equipment, facilities, and consultations services to support investigators in performing behavioral tasks. The Iowa Institute of Human Genetics provided snRNA sequencing services. Dr. Jocelyne Caboche, Sorbonne University, kindly provided AAV-ENK-cre virus. Dr. Joseph F. Lynch III, University of Iowa, helped maintain mouse lines.

