## Supplemental information for "A chromosome region linked to neurodevelopmental disorders acts in distinct neuronal circuits in males and females to control locomotor behavior"

### SUPPLEMENTARY MATERIALS AND METHODS

#### Drug treatment

For intraperitoneal injections (IP), Risperidone (Sigma-Aldrich, Product No. R3030) or SCH39166 (Tocris, Cat. No. 2299) was dissolved in 2% DMSO in physiological saline and injected at doses of 0.2 mg/kg in a volume of 5 ml/kg. Vehicle treatment received 5 ml/kg of 2% DMSO in saline. The mice were acclimatized to the monitoring chambers for one week before data collection to reduce the effect of a new environment. In second week, the mice were injected with vehicle at beginning of the dark cycle everyday (total 7 times of vehicle injection). In third week, the mice were injected with drug (Risperidone or SCH39166) at beginning of the dark cycle everyday (total 7 times of drug injection). The data was collected for two weeks following the habituation period.

### SUPPLEMENTARY FIGURE LEGENDS

Figure S1. Cell type sorting UMAP results show similar cell portions between sex and genotypes.

Figure S2. Marker gene expressions show parallel results between sex and genotypes. The size of the dots displays the percent of marker gene expression from 100% to 0%. The color of the dots presents genotype and sex; light orange (wt female), dark orange (16p11.2 del/+ female), light blue (wt male), and dark blue (16p11.2 del/+ male). We inadvertently detected several cortical neuron populations from adjacent cortex regions due to their proximity to the striatum.

Figure S3. Distinct synaptic functional changes are detected in D1-SPNs and D2-SPNs of 16p11.2 del/+ male mice. (A, C, E) Recording of synaptic events in NAc D1-SPNs of male 16p11.2 del/+ mice (n = 20 wt cells from 4 animals, n = 22 16p11.2 del/+ cells from 4 animals) show reduced half-width (p=0.0005. C) and decay time (p=0.0029, E) of the synaptic currents, but the rise-time remined unchanged (p=0.4518, A). (B, D, F) Recording of synaptic events in NAc D2-SPNs of male 16p11.2 del/+ mice (n = 15 wt cells from 5 animals, n = 18 16p11.2 del/+ cells from 3 animals) show comparable rise-time (p=0.3591, B), half-width (p=0.2598, D), and decay time (p=0.5546, F) of the synaptic currents. Error bars represent SEM. *p < 0.05.

Figure S4. The sex-specific transcriptomic changes in 16p11.2 del/+ male mice correlate with wildtype sex differences. (A, B) Quadrant plots depicting the correlation between differentially expressed genes in D1-SPNs (R=0.51, p=8.5e-08, A) or D2-SPNs (R=0.49, p=3.3e-08, B) in 16p11.2 del/+ male mice. X-axis shows DEG fold change in the D1-SPNs (A) and D2-SPNs (B) in 16p11.2 del/+ male mice compared to wt male mice. Y-axis displays fold change in wt female mice compared to wt male mice. (C, D) Quadrant plots depicting the correlation between differentially expressed genes in D1-SPNs (R=-0.56, p=0.01, A) or D2-SPNs (R=-0.33, p=0.15, B) in 16p11.2 del/+ female mice. X-axis shows DEG fold change in the D1-SPNs (A) and D2-SPNs (B) in 16p11.2

del/+ female mice compared to wt female mice. Y-axis displays fold change in wt female mice compared to wt male mice. (E) Quadrant plot depicting the correlation between differentially expressed genes identified in bulk RNA-seq of striatum in 16p11.2 del/+ mice compared to wt mice (R=0.360, p<0.001). X-axis shows DEG fold change in 16p11.2 del/+ male mice compared to wt male mice. Y-axis displays fold change in wt female mice compared to wt male mice. (F) Quadrant plot depicting the correlation between differentially expressed genes identified in bulk RNA-seq of striatum in 16p11.2 del/+ mice compared to wt mice (R=-0.082, p=0.4). Y-axis shows DEG fold change in 16p11.2 del/+ female mice compared to wt female mice. X-axis displays fold change in wt female mice compared to wt male mice. (G) Schematic depicting the expectation (upper) and our observation (lower), suggesting that the baseline sex difference correlate with DEGs in 16p11.2 del/+ male mice.

Figure S5. Locomotor behaviors are shown in double positive mice (experimental group) and other three genotype mice. The plots (A, B, C, D) display the separated control group by three genotypes in figure 4B,D,G,I, respectively.

Figure S6. Both male and female 16p11.2 del/+ mice show hyperactivity. (A) Activity monitoring data shows hyperactivity in male 16p11.2 del/+ mice compared to wild type littermates. Main effect of genotype F(1, 27)=12.75, p=0.0014; main effect of time, F(23, 621)=13.23, p<0.0001; genotype x time interaction, F(23, 621)=4.973, p<0.0001. Post hoc shows significant differences between male 16p11.2 del/+ mice and control littermates in 13-18 hour time slots. (B) Activity monitoring data shows hyperactivity in female 16p11.2 del/+ mice compared to wild type littermates. Main effect of genotype F(1, 17)=9.767, p=0.0062; main effect of time, F(23, 391)=24.35, p<0.0001; genotype x time interaction, F(23, 391)=6.944, p<0.0001. Post hoc shows significant differences between female 16p11.2 del/+ mice and control littermates in 12-22 hour time slots. Error bars represent SEM. ***p < 0.001.

Figure S7. 16p11.2 flox mice do not show locomotor activity alteration. (A) Activity monitoring data does not show difference between male 16p11.2 flox mice and wild type littermates. Main effect of genotype F(1, 21)=0.02127, p=0.8854; main effect of time, F(23, 482)=24.68, p<0.0001; genotype x time interaction, F(23, 482)=0.5081, p=0.9735. No change is reported in post hoc between male 16p11.2 flox mice and control littermates. (B) Activity monitoring data does not show difference between female 16p11.2 flox mice and wild type littermates. Main effect of genotype F(1, 19)=0.0077, p=0.9308; main effect of time, F(23, 437)=16.91, p<0.0001; genotype x time interaction, F(23, 437)=0.7952, p=0.7386. No change is reported in post hoc between female 16p11.2 flox mice and control littermates. Error bars represent SEM.

Figure S8. Risperidone (D2R antagonist, 0.2 mg/kg, IP), but not SCH39166 (D1R antagonist, 0.2 mg/kg, IP), reduces hyperactivity in male 16p11 del/+ mice. (A, B) Infrared beam breaks are plotted across the 24-hr in 1-hr bins (A) and in 24-hr (B). Activity monitoring data displays decreased hyperactivity after Risperidone injection (0.2 mg/kg, IP) compared to vehicle injection (saline, IP) in male 16p11.2 del/+ mice. Main effect of treatment F (1, 5)=4.963, p=0.0764 (A) and paired t-test shows p=0.0312 between vehicle treatment and Risperidone treatment (B). (C,D) Activity monitoring data displays similar activity after SCH39166 injection (0.2 mg/kg, IP) compared to vehicle injection (saline, IP) in male 16p11.2 del/+ mice. Main effect of treatment F (1, 7)=3.409, p=0.1073 (C) and paired t-test shows p=0.1953 between vehicle treatment and SCH39166 treatment. *p < 0.05.

Figure S9. Cre-mediated 16p11.2 del/+ is validated using *df* allele PCR. (A) Schematic of the chromosome engineering strategy of cell type-specific 16p11.2 hemideletion. Adapted from (Horev et al. 2011) (B) PCR products using primers specific for the deletion show df allele band from the striatum tissue from 16p11.2 del/+ mice. (C) PCR products using primers specific for the deletion show *df* allele band in the striatum tissue from D1-16p11.2 del/+ mice but not from control mice. (D) *df* allele band was detected only in the striatum tissue from D2-16p11.2 del/+ mice but not from control mice. (E) *df* allele band was detected in the cortex tissue from Ctx-16p11.2 del/+ mice but not from control mice.

Figure S10. ENK-cre virus-mediated 16p11.2 del/+ is validated using *df* allele PCR. (A) *df* allele band was detected only in ENK-cre virus injected striatum tissue from 16p11.2 flox mice but not from wt mice. (B) *df* allele band was detected only in ENK-cre virus injected striatum tissue from 16p11.2 flox mice but not from the cerebellum tissue.
