## Supplemental figures for "A chromosome region linked to neurodevelopmental disorders acts in distinct neuronal circuits in males and females to control locomotor behavior"

Supplementary Figure 1. Cell type sorting results show similar cell portions between sex and genotypes

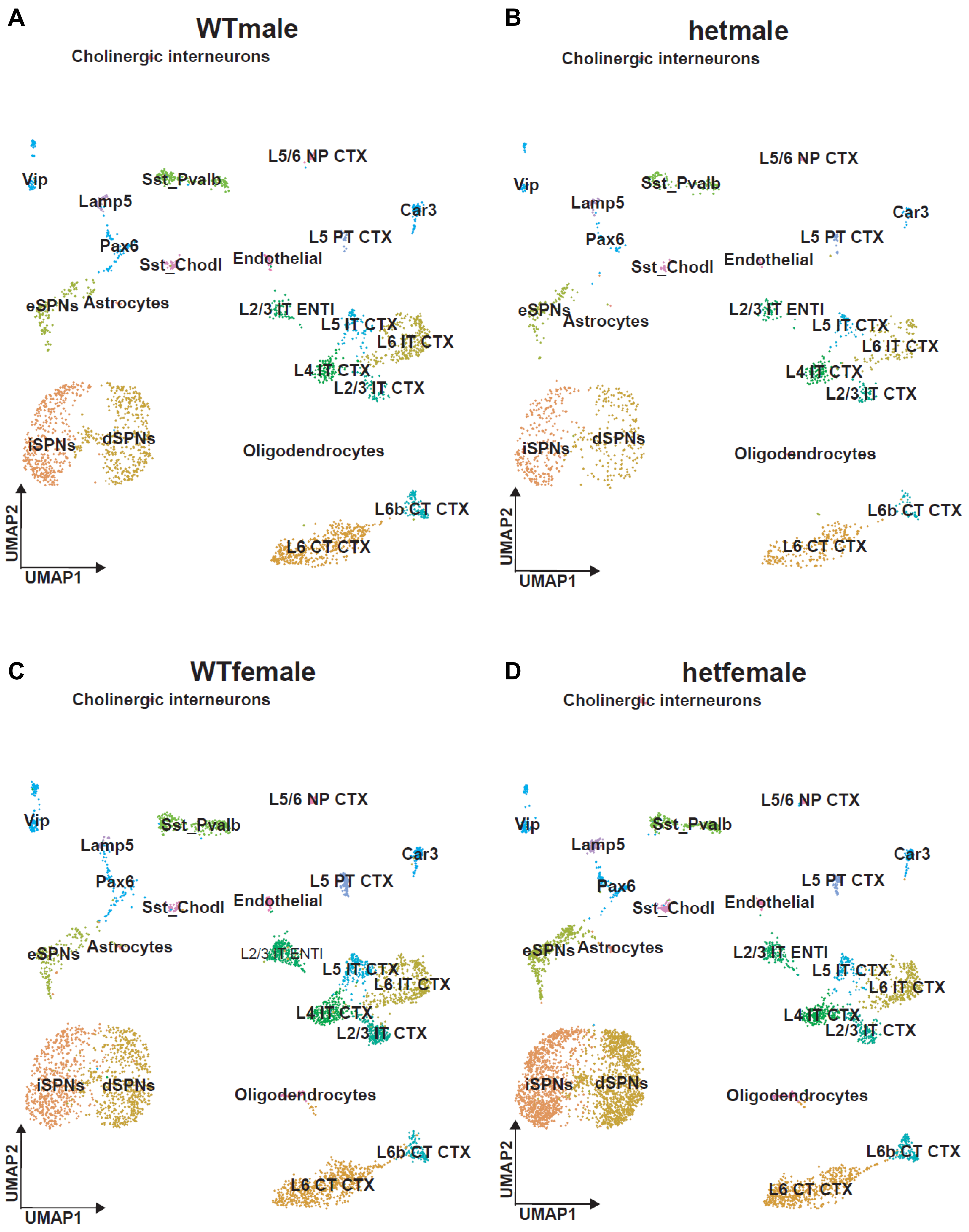

**Supplementary Figure 2. Marker gene expressions show parallel results between sex and genotypes**

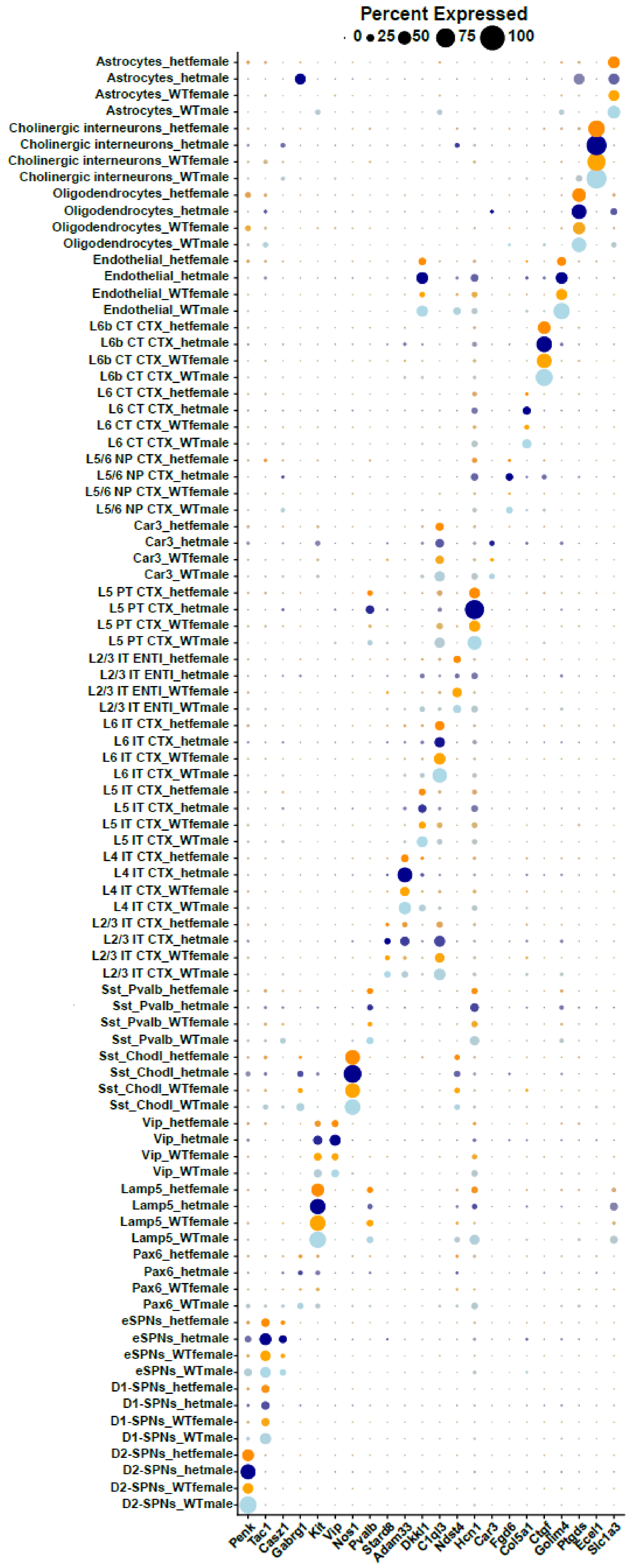

**Supplementary Figure 3. Distinct synaptic functional changes are detected in D1-SPNs and D2-SPNs of 16p11.2 del/+ male mice**

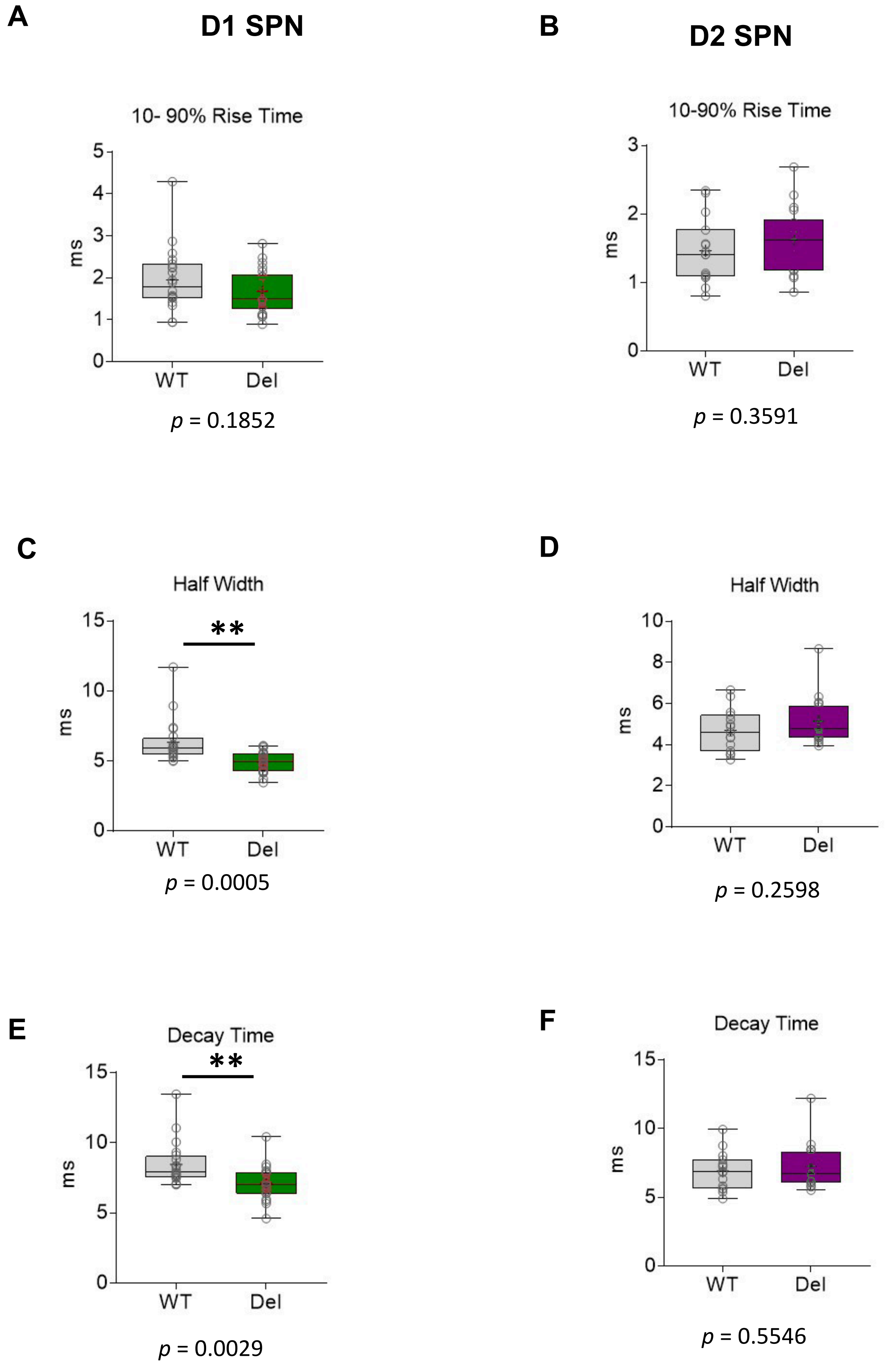

**Supplementary Figure 4. The sex-specific transcriptomic changes in 16p11.2 del/+ male mice correlate with wildtype sex differences**

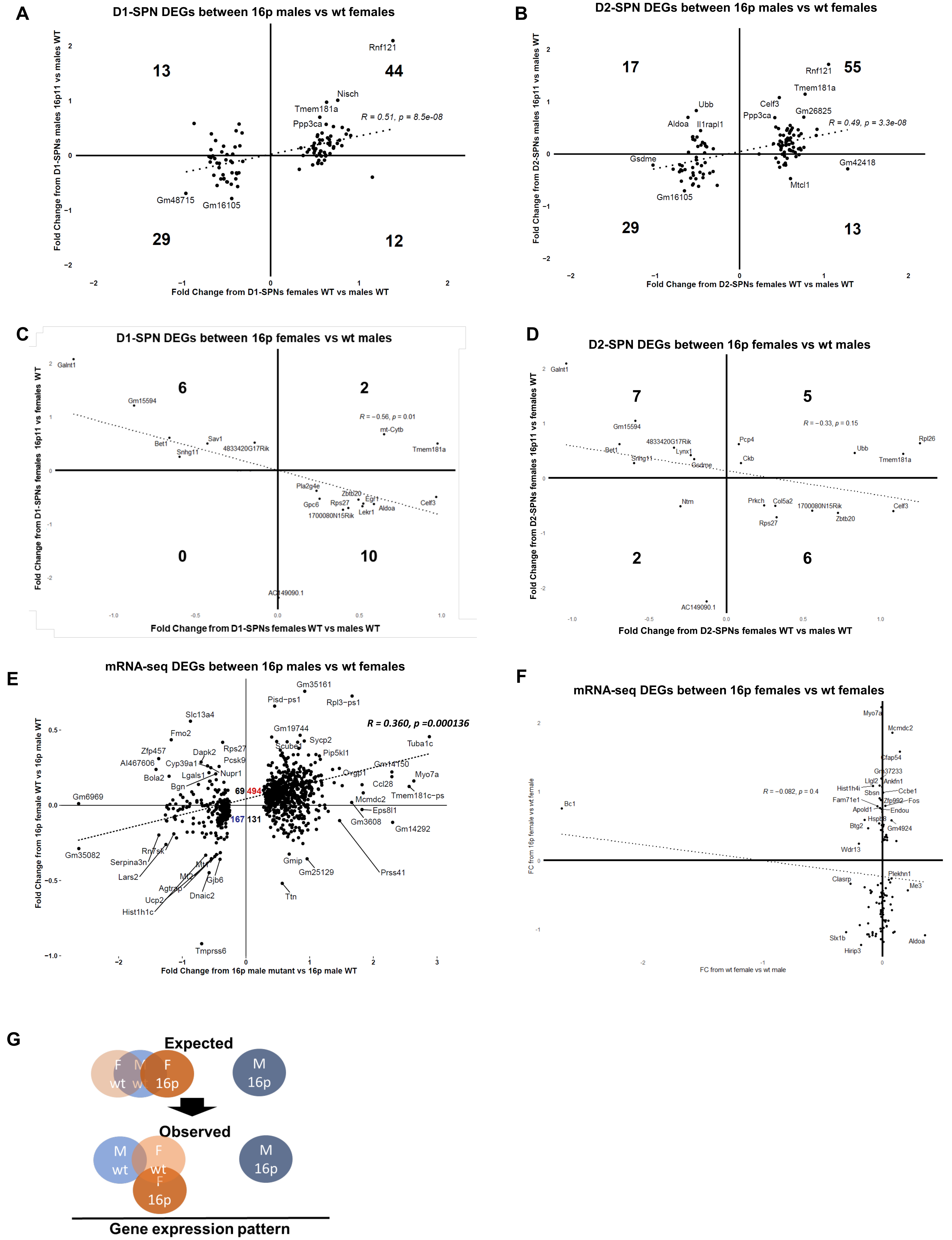

**Supplementary Figure 5. Locomotor behaviors are shown in double positive mice (experimental group) and other three genotype mice.**

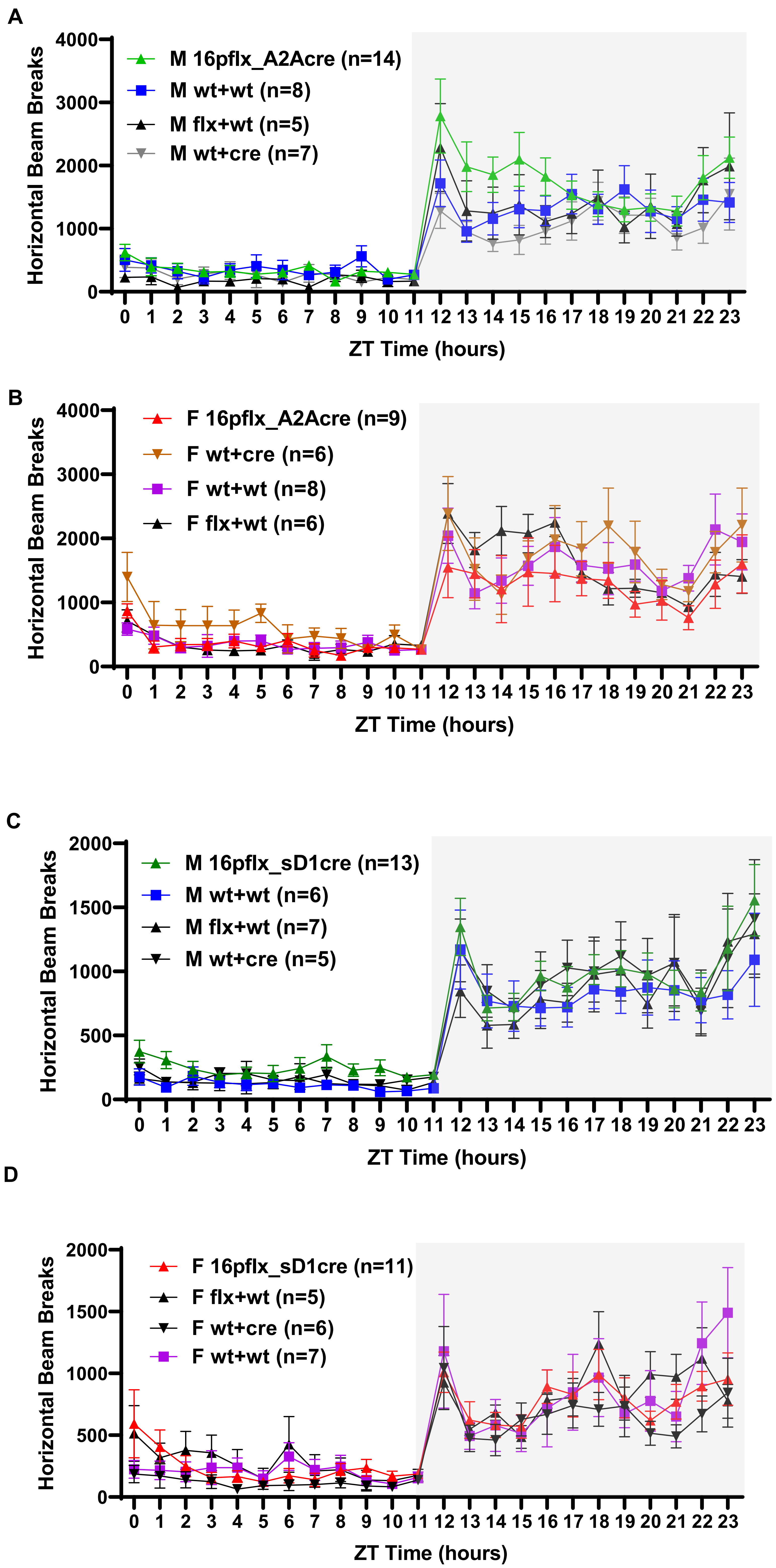

**Supplementary Figure 6. Both male and female 16p11.2 del/+ mice show hyperactivity.**

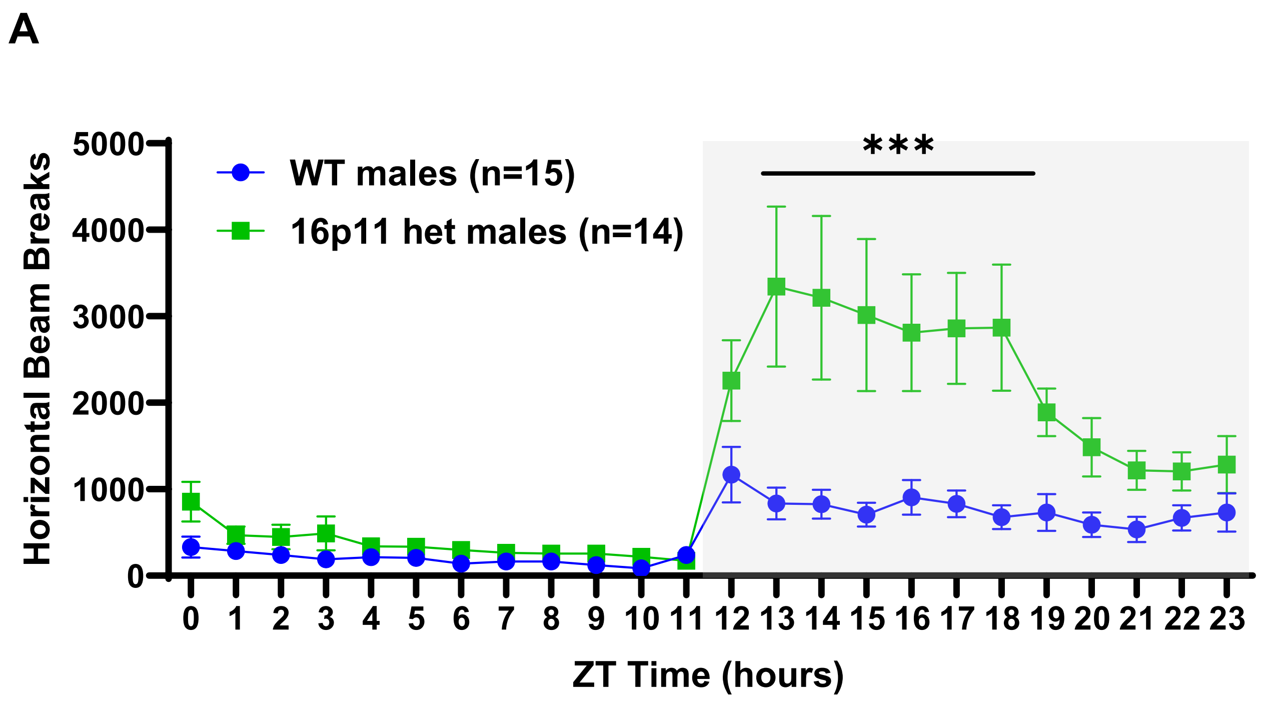


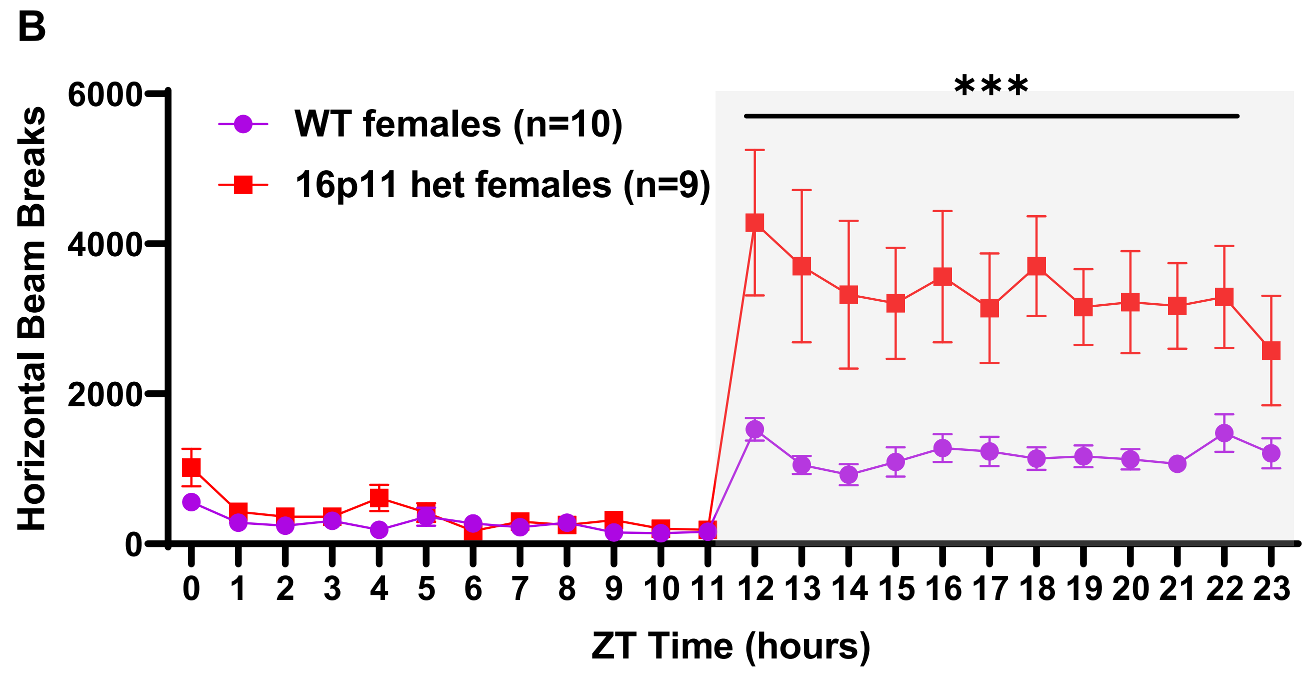

**Supplementary Figure 7. 16p flox mice do not show locomotor activity alteration.**

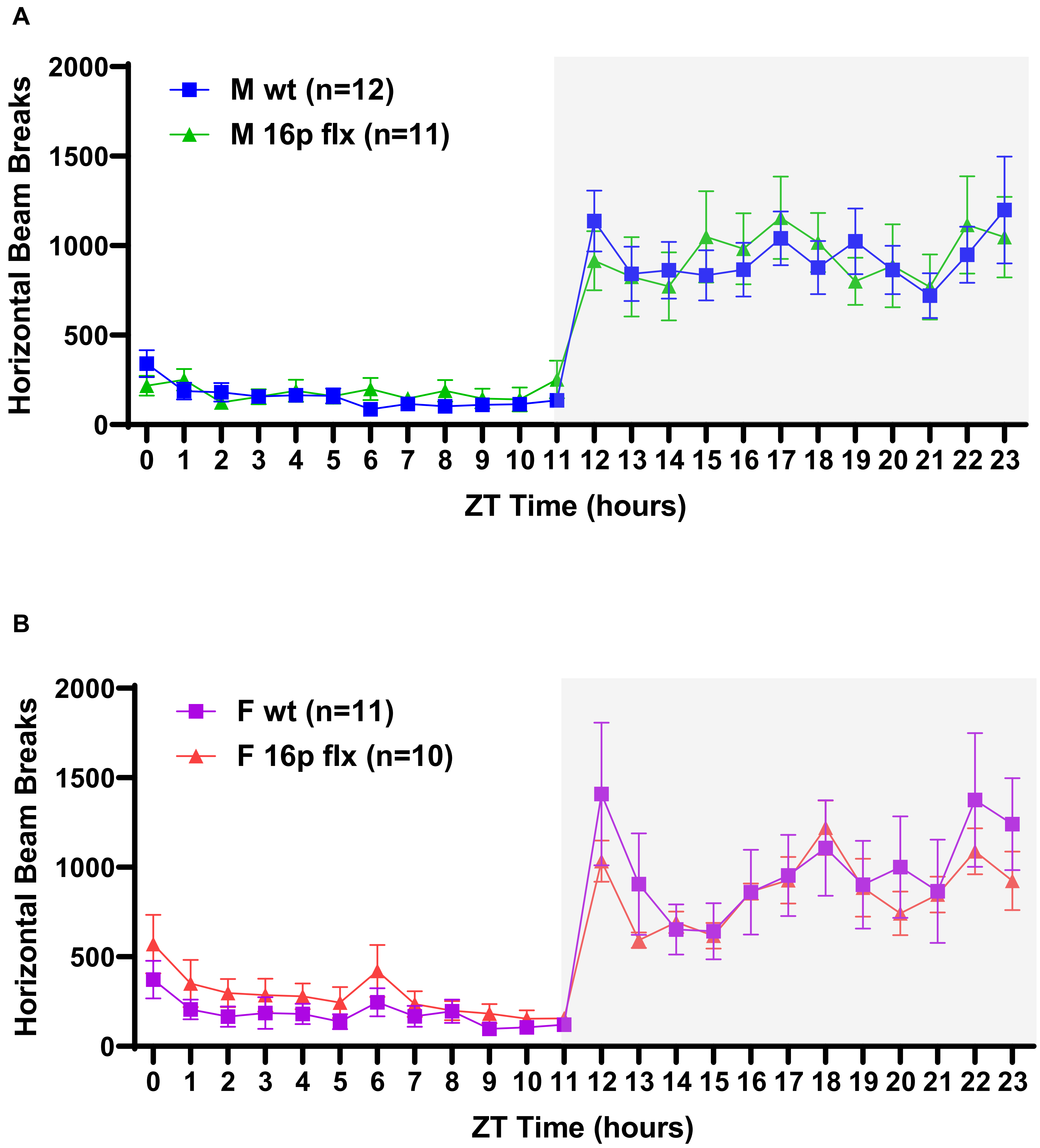

**Supplementary Figure 8. Risperidone (D2R antagonist), but not SCH39166 (D1R antagonist), reduces hyperactivity in male 16p11 del/+ mice.**

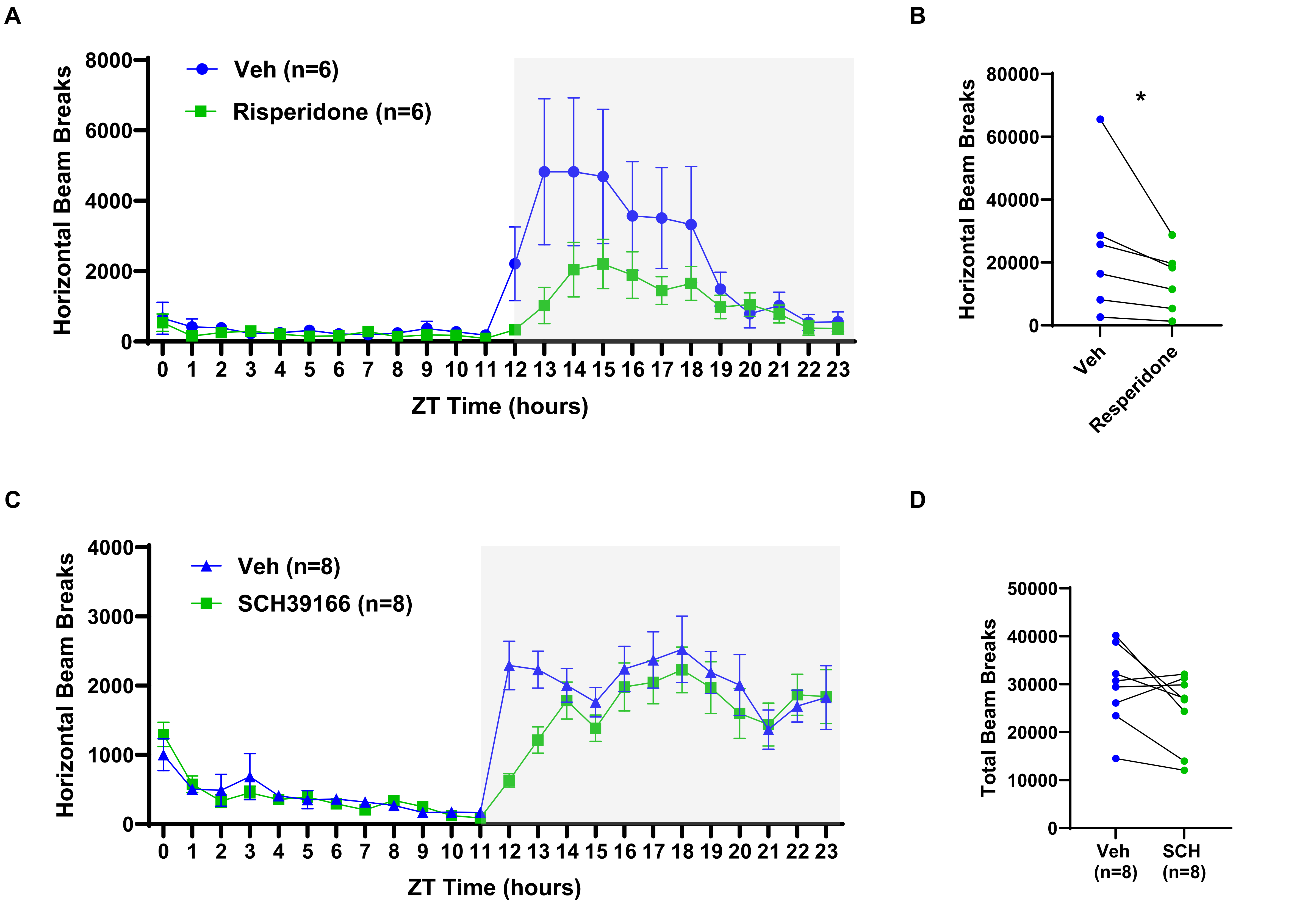

**Supplementary Figure 9. Cre-mediated 16p11.2 del/+ is validated using df allele PCR.**

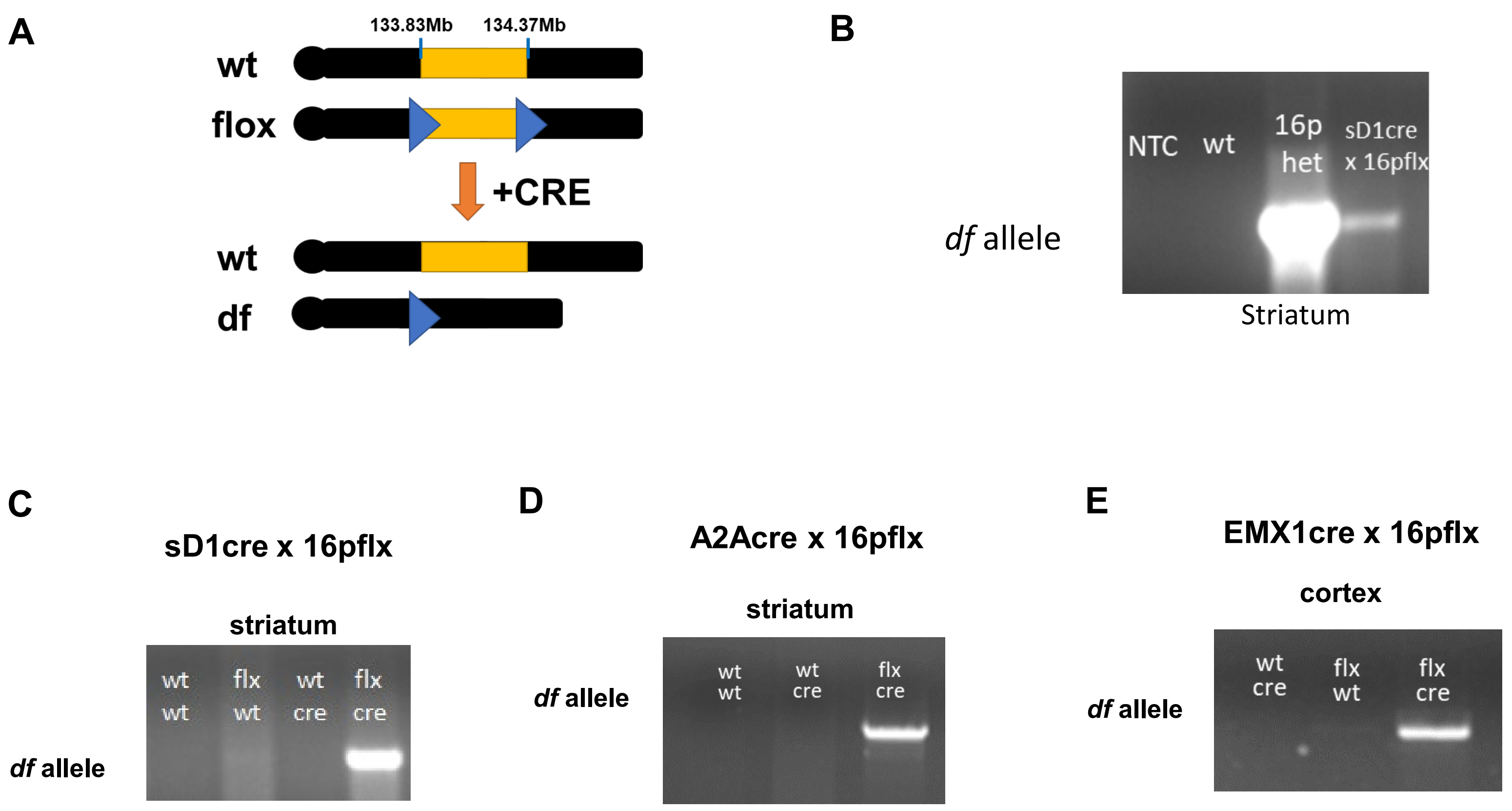

**Supplementary Figure 10. ENK-cre virus-mediated 16p11.2 del/+ is validated using df allele PCR.**

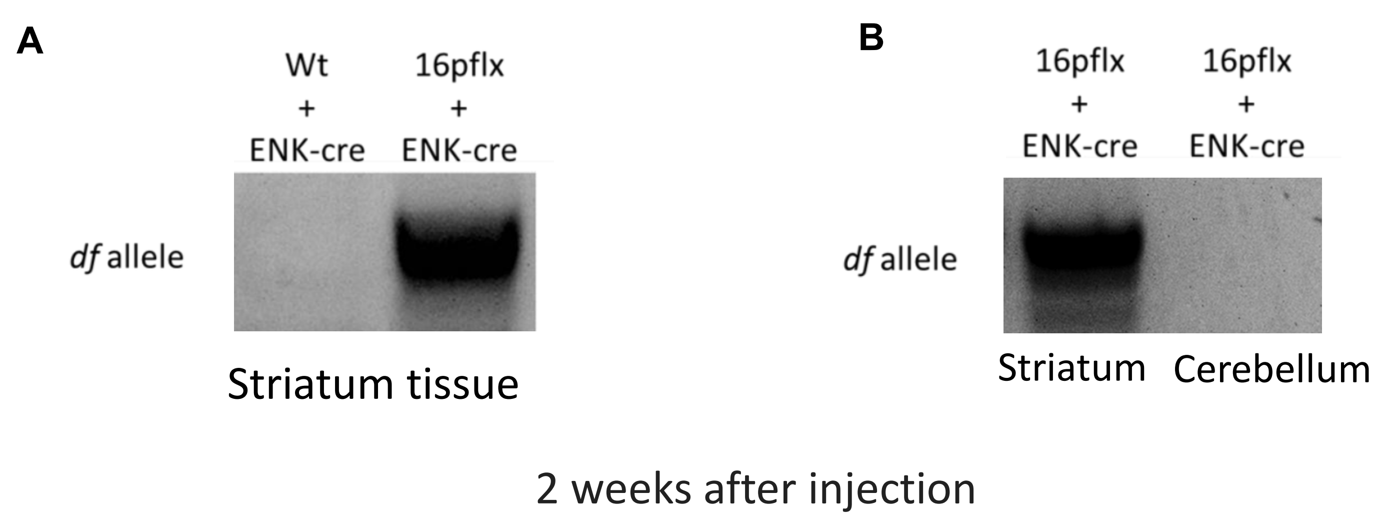
